# Engineering Cortical Networks: An Open Platform for Controlled Human Circuit Formation and Synaptic Analysis *In vitro*

**DOI:** 10.1101/2025.03.07.642064

**Authors:** Pacharaporn Suklai, Taylor F. Minckley, Cathleen Hagemann, Karolina Faber, Rosalind Norkett, Ludovica Guetta, Kelly O’toole, Bethany Geary, Michael J. Devine, Andrea Serio

## Abstract

Neuronal circuits are complex networks formed by specific neuron connections across brain regions. Understanding their development is key to studying circuit-related dysfunctions in brain diseases. Human-induced pluripotent stem cell (iPSC) models aid in this research but lack precise architecture, limiting insights into neuronal interactions and activity-dependent processes. Microfluidic technologies offer structural control but are restricted by closed systems that hinder 3D integration, scalability, and cell retrieval.

To address these limitations, we developed an open cortical network platform integrating iPSC-derived cortical neurons with bioengineering techniques. Using a polydimethylsiloxane (PDMS)-based microgroove topography and a cell plating guide, we created “neuronal nodes” for controlled circuit assembly. This design enables large-scale functional cortical circuits without physical barriers, allowing optogenetic control of neural activity and flexible network modifications, including cellular composition, neurite directionality, and synapse formation.

The open design facilitates neuronal material accessibility, supporting multi-level analyses such as proteomics. This platform serves as a powerful tool for investigating neuronal network development and function, offering new opportunities to study both normal and pathological states, including molecular changes linked to connectivity loss in brain diseases.

## 2. Introduction

Advances in neuroscience have long been driven by the development of innovative tools and models that facilitate the study of human brain function and dysfunction. Among these, the discovery of human induced pluripotent stem cells (iPSCs) by Takahashi and colleagues in 2007^1^ marked a turning point, enabling unprecedented opportunities to investigate complex human biology through patient-derived cellular systems. iPSCs offer a transformative platform for modeling human diseases^2^, drug discovery, and regenerative therapies^3^ and address a critical gap in neuroscience research. Thanks to their application we have now human-specific cellular *in vitro* models that complement the limitations of animal models, as they often fail to fully capture the molecular, functional, and cellular nuances of human neurological disorders.

Particularly in the study of neuronal circuit assembly, for either basic neurobiology or disease modelling, *in vitro* human models are of crucial importance. For example, modelling the human cerebral cortex, a region essential for higher-order functions such as cognition, memory, and sensory processing, present several challenges as rodents and other model species present fundamental differences in cellular composition, architecture, and developmental trajectories fail to replicate the full spectrum of human cortical function and also disease progression. By leveraging iPSC-derived neurons, researchers can construct patient-specific cortical models to investigate the cellular and molecular processes underlying neurological disorders with greater fidelity.

Conventional *in vitro* neuronal culture techniques have significantly advanced our understanding of basic neuronal functions and synaptic connectivity. Cultured neurons derived from iPSCs recapitulate many key features of *in vivo* neurodevelopment, including dendritic and axonal growth, synaptic transmission, and network activity. These 2D cultures provide a simplified and controlled environment for studying neuronal function, offering insights into action potential generation, synaptic plasticity, and circuit dynamics. However, despite these advantages, conventional 2D models have limitations when it comes to modeling the complexity of *in vivo* neural circuits. Randomized orientation and interactions in 2D cultures often result in reduced functional relevance, particularly when investigating organized and activity-dependent neural processes^4^. To address these limitations, bioengineering techniques have been employed to impose structure and enhance circuit functionality.

Microfluidic platforms represent a pivotal advancement in neuronal circuit modeling, offering precise control over cell placement, connectivity, and directional growth. These devices utilize compartmentalized designs to isolate neuronal compartments, such as soma and axons, enabling targeted studies on synaptic connectivity and axonal transport. The incorporation of microchannels and synaptic compartments allows researchers to manipulate neuronal interactions and observe bidirectional or unidirectional connectivity patterns^5,6^. For example, microfluidic systems have been used to construct cortical-thalamic and cortico-striatal circuits, facilitating studies on synaptic plasticity, network activity, and disease-specific mechanisms such as those seen in Huntington’s disease^7^. Moreover, microfluidic platforms support synaptic competition models^8^ and the precise formation of unidirectional networks^9^, mimicking the connectivity observed in the human brain.

Despite their versatility, microfluidic systems face challenges in scalability, integration with 3D cultures, and accessibility for certain experimental techniques. These limitations have driven the development of more complex 3D neuronal models, such as brain organoids and assembloids^10^, which can recapitulate to different extent the connectivity of given circuits in a developmentally relevant fashion.

However, organoids and other 3D cultures also have inherent limitations: for example, their structure complicates the recording and manipulation of neuronal activity, and their complexity makes it difficult to deconstruct and analyze individual circuit components.

These considerations highlight the need for a new *in vitro* platform that bridges the gaps in current models. A scalable and reproducible platform capable of reconstructing and manipulating connectivity, as well as recovering circuit materials for molecular studies could significantly expand the scope of research questions and provide deeper insights into cortical network function. For example, analyzing cellular and molecular changes during specific network events may help link molecular alterations to processes underlying network formation and function.

The present study aims to address these challenges by developing a new cortical circuit platform called BIOCONNET (BIOengineered COrtical Neuronal NETwork). This open-system platform seeks to integrate the strengths of 2D and 3D neuronal models while overcoming their limitations. BIOCONNET enables precise manipulation of circuit components, supports scalable analyses such as transcriptomics and proteomics, and provides a versatile framework for studying the intricacies of cortical network formation and function. By combining bioengineering and iPSC-derived neurons, this platform represents a significant step forward in the modeling of human cortical circuits, offering new opportunities for advancing our understanding of brain health and disease.

## 3. Results

### Assembly of a Barrier-Free Platform using Microtopography and Plating Devices resulting in Controllable Cell Positioning and Neurite Directionality

To create an open platform without obstructing barriers from the top – meaning no part of the neurons is enclosed in PDMS or unretrievable - while maintaining the control over the patterns of neuronal culture, we introduced a PDMS-based microtopographic surface combined with a plating device.

Previously, we developed bioengineered devices composed of two main components in order to generate a cell culture platform with controllable cell positioning and neurite directionality^11^. The first component consists of PDMS microgrooves with 10 µm × 10 µm (width x height), created via soft lithography on a silicon master microfabricated through photolithography (**Figure 1A**). These micropatterned surfaces are biofunctionalized with O^2^ plasma treatment followed by poly-D-lysine (PDL) and laminin coating, allowing for cell adhesion. Cortical neuron progenitors derived from iPSCs^12^ are then plated on this biofunctionalized PDMS surface and differentiated into mature neurons (see **Supplementary Figure 1** for the differentiation process, along with characterization confirming their neuronal identity and functionality). We observed that cortical neurons on the micropattern aligned along the groove direction, in contrast to the lack of specific directionality on flat surface (**Figure 1B**). In addition, MAP2-labled dendrites branched beyond the grooves, enabling network formation between neurons located in different grooves. This microgroove array provides an organization of neurites while retaining freedom of network formation.

**Figure 1.**
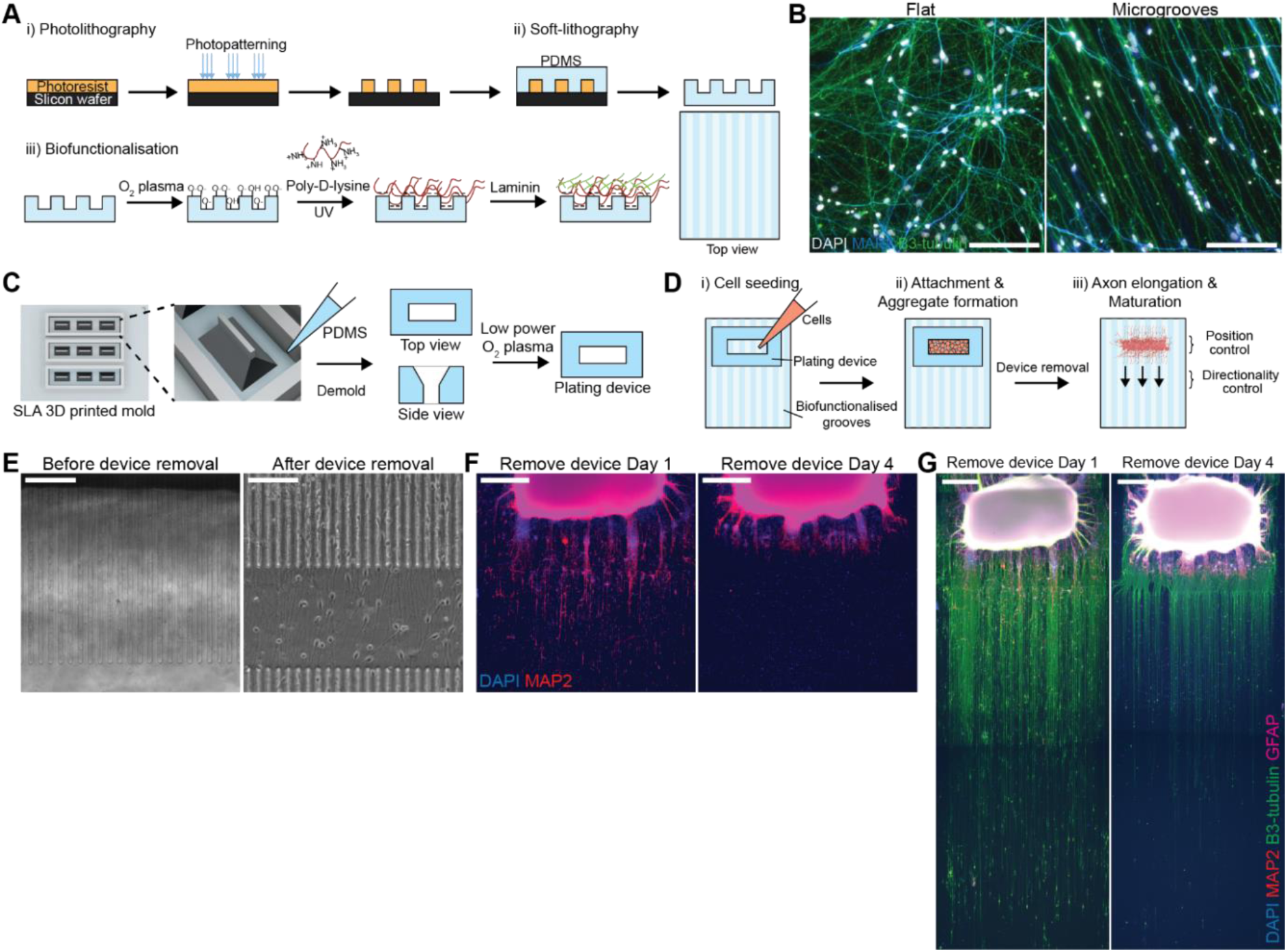
An open platform with controllable cell location and neurite directionality. A) Schematic overview of the production of micropatterned substrate for neurite directional control. B) Representative images of cortical neurons plated on flat and microgrooved surfaces, in which cells were terminally differentiated for 14 days. Scale bar: 100 μm. C) Schematic overview of the generation process of PDMS plating devices for cell location control. D) Schematic overview of the cell seeding process to create neuronal aggregates with neurites projecting outside and aligning with microgroove patterns. E) Representative images of cellular confinement upon the temporarily sealing between the plating device and microgroove substrate, creating microchannels that allow neurites to pass through (left), and once the device was removed the open system allows neurons to move freely (right). Scale bar: 100 μm. F) and G) Representative images of neuronal aggregates generated with device in place for 1 day (left) and 4 days (right) before removal and followed by fixation and staining after 7 days of culture. Scale bar: 500 μm.

**Supplementary Figure 1.**
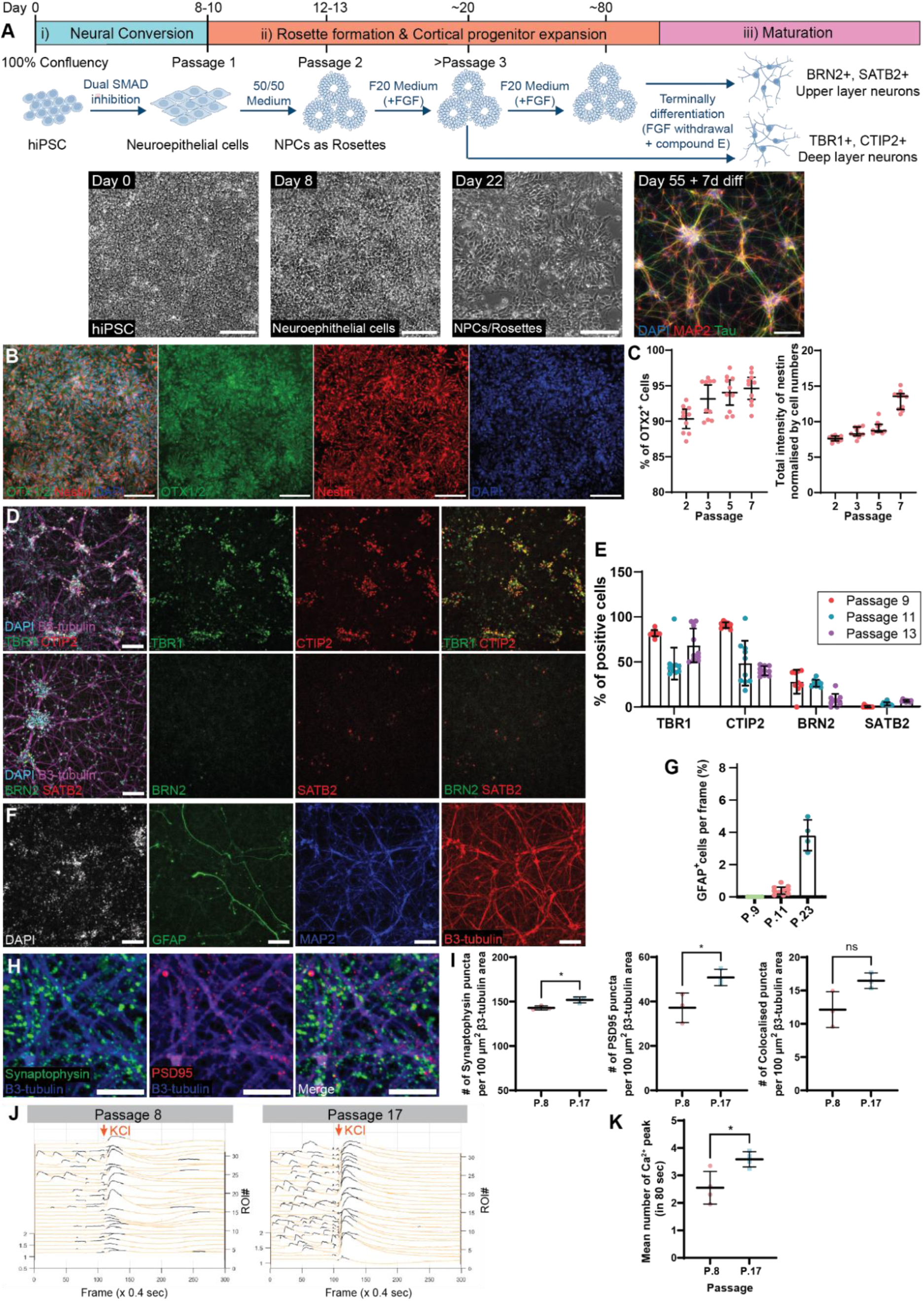
Generation and characterization of iPSC-derived cortical neurons. **A**) Schematic overview of cortical neuron induction and differentiation from iPSCs, containing 3 main phases: i) neural conversion, ii) neural rosette formation and proliferation of cortical neuron progenitors, and iii) terminal differentiation and maturation. Representative images show the progression from iPSCs (Day 0) to neuroepithelial cells (Day 8), followed by the formation of neural progenitor cells organized into ‘rosette’ structures (Day 22). Upon FGF withdrawal, these progenitors terminally differentiate into cortical neurons (Day 55 + 7 days). Staining includes DAPI for nuclei, MAP2 for dendrites, and Tau for axons. Scale bar: 100 μm. **B**) Representative images of cortical neuron progenitors expressing OTX1/2, a forebrain cortical progenitor marker, and Nestin, a general neural marker. Scale bar: 100 μm. **C**) Quantification of OTX1/2 positive cells (left) and Nestin expression, presented as total Nestin intensity normalized to the number of cells (right) in cortical neuron progenitor cultures across different passages. Data points represent each field of view (FOV) from 2 technical replicates, with results shown as the median ± interquartile range. **D**) Representative images of cortical neurons at passage 13, terminally differentiated for 7 days, stained for cortical neuron subtype transcription factors: TBR1 and CTIP2 (deeper layer) and BRN2 and SATB2 (upper layer). Scale bar: 100 μm. **E**) Percentage of cells positive for each transcription marker across different passages. Data points represent each FOV from 2 technical replicates, with results shown as mean ± S.D. **F**) Representative images of cortical neuron culture at passage 11, terminally differentiated for 7 days, stained for astrocyte marker GFAP, and neuronal markers MAP2 and β3-tubulin. Scale bar: 100 μm. **G**) Percentage of GFAP positive cells in culture across different passages. Data points represent each FOV from 2 technical replicates, with results shown as mean ± S.D. **H**) Representative images of cortical neurons from passage 17 (day 70), showing the expression of synaptic proteins synaptophysin and PSD95, with β3-tubulin as a neuronal cytoskeleton marker. Scale bar = 10 µm. **I**) Quantification of synaptic puncta: pre-synaptic puncta (synaptophysin, left), post-synaptic puncta (PSD95, middle), and colocalised puncta (right) per 100 μm^2^ β3-tubulin. Data represent well averages from 1 independent experimental block with 1 cell line each with 3 technical replicates. Synaptophysin data are expressed as median ± interquartile range and analysed using the Mann-Whitney test, while PSD95 and colocalized puncta are expressed as mean ± SD, and analysed using an unpaired t-test (* P<0.05, ns: non-significant). **J**) Representative calcium spike traces from individual regions of interest (ROIs) drawn around the soma of cortical neurons, terminally differentiated for 14 days from progenitors at passage 8 (day 38) and passage 17 (day 70). Neurons were stimulated with 30 mM KCl at an indicated arrow. **K**) Quantification of the total number of calcium spikes at baseline over an 80-second period. Data represent well averages from 1 independent experimental block with 1 cell line, each with 4 technical replicates. Results are expressed as mean ± SD, using unpaired t-test to test for significance (* P<0.05, ns: non-significant).

To be able to control the position of neurons in the culture, we applied the second component serving as a cell plating guide, generated by soft lithography of PDMS from stereolithography (SLA) 3D printed mould (**Figure 1C**). The plating device^11^ features an open channel with a funnel-like shape that directs cells to specific regions on the topographic substrate and promotes initial cell aggregation. Prior to cell seeding, the plating device undergoes a brief oxygen plasma treatment to enhance PDMS hydrophilicity, which facilitates dispersion of liquid and cells during seeding. To place neurons at specific positions, the plating device is placed on the microgroove substrate at the desired location, generating temporary wells into which cells are pipetted (**Figure 1D**). Once the neurons have adhered and formed aggregates, the plating device is removed, creating a fully open culture system without physical barriers (**Figure 1E**). These aggregates anchor neurons in place while allowing axons to extend outward during differentiation and maturation.

To optimize aggregate formation and minimize axonal damage from the plating device removal, we adjusted the culture period within the plating device. Cortical neuronal aggregates formed within a day, but some neurons appeared to migrate around the aggregate after device removal (**Figure 1E**). Extending the culture time within the device reduced migration but also limited axon growth (**Figure 1F, G**). We determined that a 48-hour culture period with the plating device balances cell aggregation with minimized interference in axonal growth. Our platform thus enables an open iPSC-derived cortical neuron culture system with precise cell positioning and neurite orientation. Unlike enclosed microfluidic systems, this open design offers flexible scalability, customization for neuronal arrays, and compatibility with 3D cultures, facilitating broader applications in neuroscience research.

### Critical Cell Number for Single, Compact Neuronal Node Formation in Bioengineered Arrays

Before establishing a controlled array of cortical circuits, we first aimed to create a ‘neuronal node’ — a confined group of cortical neurons that receive similar synaptic inputs and can function as a unit within a circuit. To achieve this, we used our optimized bioengineered device to create a cortical aggregate that would serve as this node. However, inconsistent aggregate formation was observed, as *in vitro* cortical neurons exhibit spontaneous self-organization, typically forming small clusters typically 180-250 μm in diameter, even when cultured using the plating device (**Figure 2A**). We then explored conditions that consistently yield uniform nodes optimal for circuit assembly, focusing on designs that closely matched the self-organized cluster size, while allowing practical plating and maximal neurite exposure for effective circuit connectivity. This led us to design a long, rectangular-shaped node. To optimize this node configuration, we evaluated two key parameters: the device width, which could help guide the cluster into a linear formation, and the cell number, increasing it until it filled any empty spaces left by self-organization to achieve uniform aggregation (**Figure 2B**).

**Figure 2.**
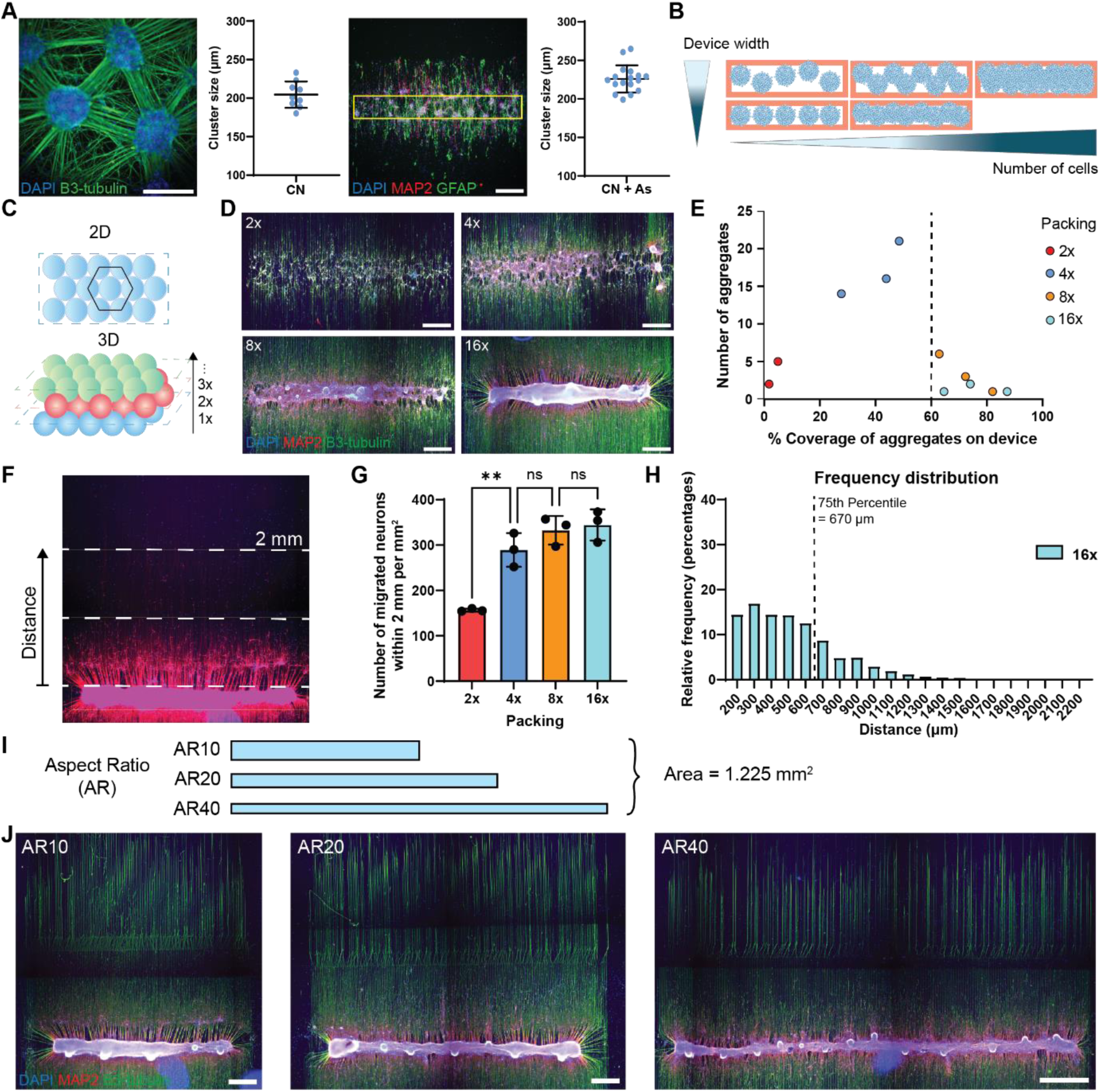
Neuronal node formation based on neuronal self-organising size requires a threshold number of cells for neuronal node packing and coherence. A) Representative images of cortical neurons forming small clusters with their size quantification from 2D conventional culture (left), scale bar: 200 μm, and when seeding using plating devices and astrocytes (right), scale bar: 500 μm. Data represents the average fields of view (FOVs) from 3 biological replicates, each with 3 well replicates for 2D samples, and 17 technical replicates across 3 rectangular device sizes for samples seeded with plating devices. B) Schematic overview of the hypothesis to generate neuronal node based on spontaneously formed clusters (blue spheres) that reduced plating device width will arrange small clusters into single lines and increased cell numbers will fill the gap between each cluster, resulting in a compact single node. C) Schematic overview of hexagonal packing principle used for calculation of cell numbers for seeding in each device. The hexagonal packing of spheres in 2D is referred to as 1x (top) and increased by adding new layers into a 3D stack, referred as 2x, 3x, … nx (bottom). D) Aggregate formation of cortical neurons by seeding with cell numbers for 2x, 4x, 8x, and 16x hexagonal packing and after maintained in differentiation for 7 days. Scale bar: 500 μm. E) Relationship between the number of aggregate counts and the percentage coverage of total aggregates occupied on the plating area. Dash line shows a threshold number of cells required for node formation. Data points represent the measurement from an individual node from 1 independent experimental block with 1 cell line each with 3 technical replicates. F) Representative image of cortical neuron migration from the node and the illustration of the quantification area up to 2 mm in distance from the node. G) The number of migrated neurons per mm^2^ from aggregates with packing 2x, 4x, 8x, and 16x. Data points represent the measurement from an individual node, expressed as mean ± SD, one-way ANOVA test with Tukey’s multiple comparisons to test significance, **P <0.01, calculated on 1 independent experimental block with 1 cell lines each with 3 technical replicates. H) The distribution of migrated neurons from 16x aggregates throughout the 2 mm distance from the node. I) Schematic diagram of plating devices with aspect ratio (AR) 10, 20, and 40, while remaining the same plating area of 1.225 mm^2^. J) Representative images of cortical neurons plated in the plating device with AR10, 20, and 40, and after maintained in differentiation for 7 days. Scale bar: 500 μm for AR10 and AR20; 870 μm for AR40.

To systematically evaluate the required cell number for coherent node formation, we applied hexagonal packing principles^13,14^, which are theoretically the most efficient way to fit spherical objects into a box, to determine the number of cells that can be packed in a defined rectangular plating device (**Figure 2C**). Here, dissociated iPSC-derived cortical neuron progenitors approximately 15 μm in diameter, that would pack into a monolayer, are defined as 1x and progressively added layers to create a 3D structure. Cortical neuron progenitors at packings 2x, 4x, 8x, and 16x were seeded within plating devices for aggregate formation for 48 hours followed by an additional 5 days after device removal. Results showed that aggregate formation was achieved gradually with increased cell number. However, only at high numbers did the self-organised clusters become collectively coherent enough to form a single aggregate (**Figure 2D, E**). Specifically, the cell numbers with packing 2x and 4x were insufficient to form cohesive aggregates, while 8x packing began to form single large aggregates, and fully coalesced at 16x. These findings suggest that forming a stable aggregate that aligns with the device geometry requires a threshold cell number at least 8x hexagonal packing.

Since the platform is barrier-free, neuron aggregates remain dynamic following device removal, with some neurons dispersing from the initial aggregate. To assess the stability of these aggregates across different seeding densities, we analysed the migratory profiles of MAP2-positive neurons that had migrated outside the plating area, extending up to 2 millimeters (**Figure 2F**). The number of migrating neurons was proportional to the initial seeding cell numbers, with significantly higher migration observed when a cohesive aggregate was not formed. Once single, cohesive aggregates were established, migration levels remained stable, regardless of further increases in seeding cell numbers (**Figure 2G**). To further evaluate the containment of neurons within a specific area, we measured the migration distance of neurons outside the plating area. Most neurons were located within 300 μm from the plating area and 75% migrated within 670 μm, suggesting the minimum distance between nodes when constructing cortical circuits (**Figure 2H**). These findings indicate that neuron aggregation using the plating device can effectively confine neuron position within a limited area, offering a promising approach for creating neuronal nodes for circuit assembly.

To evaluate the effect of device width on guiding clustered neurons into a linear shape, we designed plating devices with different widths including 350 μm, 248 μm, and 175 μm while keeping the plating are constant and thus varying the aspect ratios to 10, 20, and 40 (**Figure 2I**). Linear neuronal node formation was achieved across all conditions, with nodes displaying small bulges along their lengths. These bulges likely represent self-organised clusters, suggesting that the reduced width of the nodes effectively guides multiple small clusters into an elongated linear arrangement (**Figure 2J**). Using these plating devices, we successfully created neuronal nodes up to a maximum length of 7 millimeters. In addition, creating longer nodes benefits the system by maximizing the projection of neurites outside the node, potentially enhancing connectivity, even when using equal cellular densities. However, as node length increased, the handling of plating became more difficult as the device removal process can disrupt aggregate attachment, particularly in longer shapes. Therefore, we chose the intermediate length as the most effective shape and efficient handling for circuit construction.

In conclusion, we identified the optimal seeding cell numbers and aspect ratio for single, compact neuronal node formation within bioengineered arrays. We chose a cell number corresponding to 16x packing, combined with an aspect ratio 20 to be the optimal condition for consistent node formation in the development of our platform.

### Astrocyte Integration and Its Effects on Node Shape Stability, Cell-Substrate Attachment, and Neuron Functionality

One aim of constructing neuronal nodes is to create nodes that maintain their shape and attachment to substrates over time, allowing for neuron maturation and circuit formation. Astrocytes are known to support the central nervous system by producing extracellular matrix (ECM) components essential for neuron stability and function^15^. To investigate the effect of astrocytes on neuronal node stability within our system, we integrated iPSC-derived astrocyte progenitors into cortical neuron progenitor mixtures for node formation (**Figure 3A**). Without astrocytes, cortical neuron-only nodes (CN-only) exhibited densely packed aggregation and contraction particularly at their ends, causing axons to lift from the substrate, resulting in axons hanging around the node (**Figure 3B**). In contrast, neuronal nodes with astrocytes (CN+As) did not display this contraction. To assess the node attachment, we measured the coverage of nodes relative to the plating area as a proxy for attachment. Although adding astrocytes increased the total cell number per node compared to CN-only node at the same packing, the observation that 16x CN-only nodes contained 1.6 times more cells than 8x CN+As nodes yet exhibited reduced substrate coverage, suggests astrocytes significantly enhance node attachment and stability (**Figure 3B, C**).

**Figure 3.**
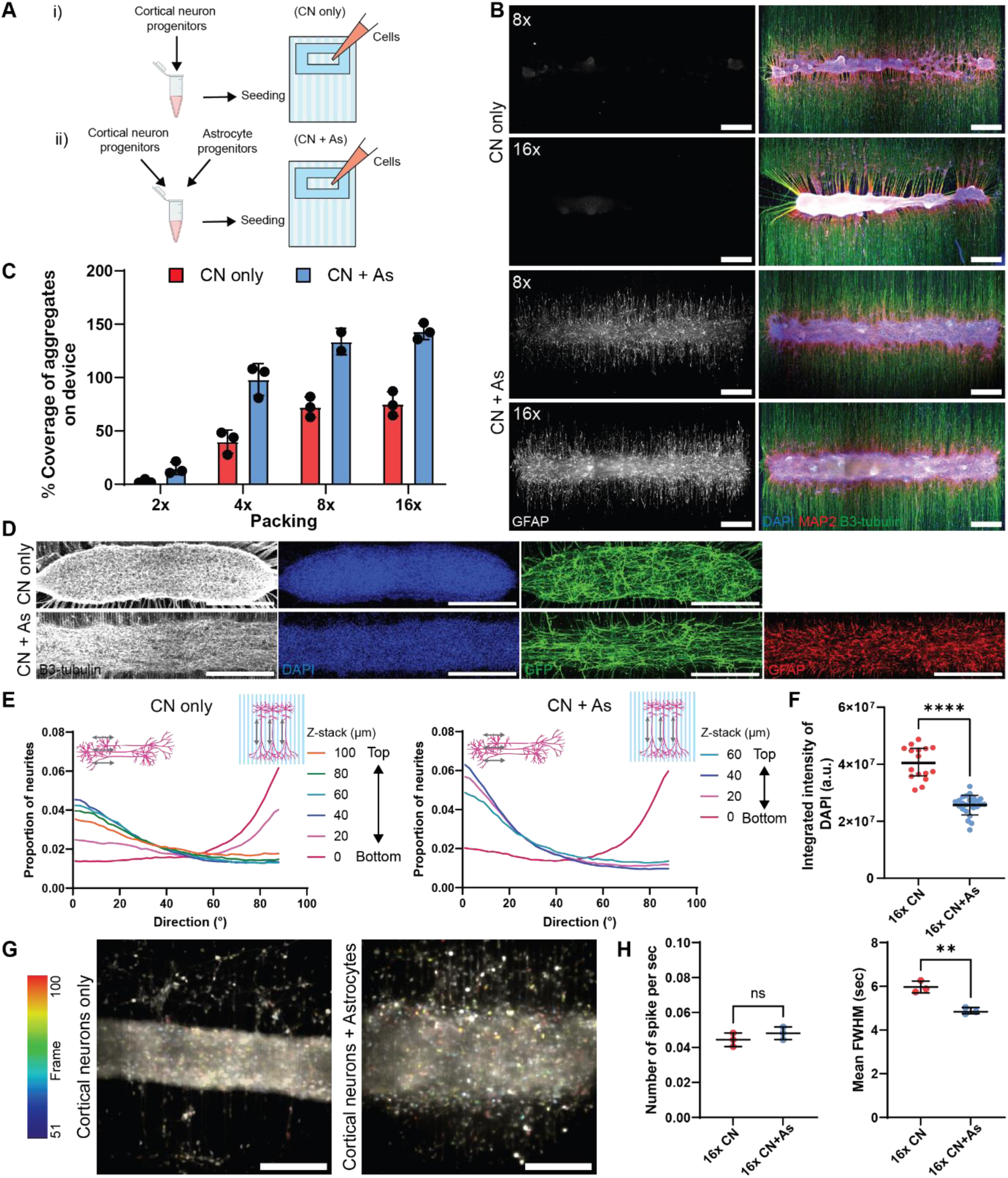
Integration of astrocytes into neuronal nodes enhances cell-substrate attachment, node shape integrity, and cortical neuron functionality. A) Schematic diagram of cell seeding conditions without astrocytes (i) and with astrocytes (ii). B) Representative images of neuronal node seeded with or without astrocytes at different cell numbers. Scale bar: 500 μm. C) Percentage area coverage of aggregates occupied on the device area. Data points represent the measurement from an individual node. D) Representative images of cellular components inside the node. Sparse GFP transfected cortical neurons show the alignment of neurites inside the node. Scale bar: 500 μm. E) Directionality of GFP positive neurites across z-stack of the node starting from PDMS surface at 0 μm to the top plane, where 0° refers to longitudinal side of the node and 90° refer to perpendicular direction to the node. Analyses were performed on 1 independent experimental block with 1 cell line each with 2-3 technical replicates. F) Node density based on integrated intensity of DAPI. Data represents the averages of 3 middle slices of the node, with 8 FOVs per node from 1 independent experimental block with 1 cell line each with 2-3 technical replicates, expressed as mean ± SD, unpaired t test, ****P<0.0001. G) Temporal colour-coding images of neuronal spikes indicated by Fluo-4AM. Scale bar: 200 μm. H) Neuronal activity analysis showing the number of calcium spikes per second (left) and full width half maximum (FWHM) (right). Data represents the averages of 3 FOVs per node from 1 independent experimental block with 1 cell line each with 3 technical replicates, expressed as mean ± SD, unpaired t test, ** P<0.01, ns: non-significant.

To further investigate how astrocytes influence the node internal structure, we created nodes with sparsely transfected GFP cortical neurons to observe neuronal organization in addition or lack of astrocytes (**Figure 3D**). The GFP-expressing neurites were double stained with GFP antibody to enhance its signal and imaged using a confocal system to analyze neurite directionality inside the nodes. In both conditions, neurites on the bottom plane, where they adhered to the substrate, aligned along the grooves (90°). However, in CN+As nodes upper-plane neurites showed a stronger horizontal alignment (0°), following the node arrangement, while CN-only nodes exhibited more diverse directionality (**Figure 3E, Supplement Figure 2**). This suggests that astrocytes improve the internal organization of neurons towards specific directions within the node. In addition, the density of cells within CN+As nodes was significantly lower than in CN-only nodes, indicating that astrocytes create a looser cellular packing that reduces node contraction (**Figure 3F**). This effect may be due to astrocyte processes penetrating the node and secreting ECM components, which adjust spacing between cell nuclei and stabilize node shape.

**Supplementary Figure 2.**
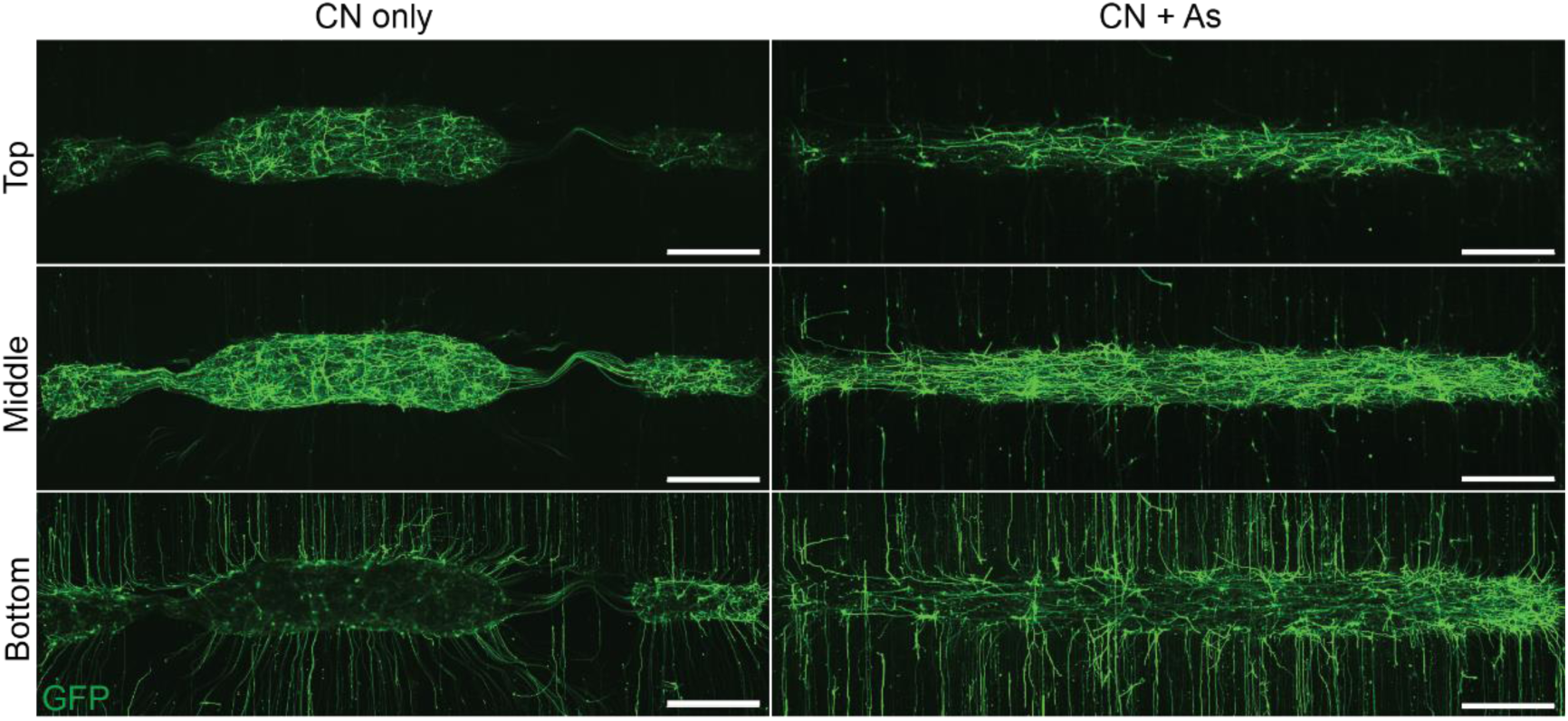
Internal organization of cortical neuron neurites. Representative images of neuronal nodes with sparse GFP cortical neurons in conditions with and without astrocytes at the top, middle, and bottom focal planes. GFP shows the alignment of neurites inside the nodes and on the topography. Scale bar: 500 μm.

In addition, astrocytes play an essential role in neural circuit development by regulating the extracellular environment that influences synapse formation and maturation^16^. Integrating astrocytes into neuronal nodes not only improved physical attachment but also enhanced neuronal circuit maturation. To investigate the effect of astrocytes on neuronal function, we measured neuronal activity within neuronal nodes formed with and without astrocytes after 14 days of differentiation. Both CN-only and CN+As nodes exhibited spontaneous calcium activity, indicating that these cells were functioning and undergoing maturation process (**Figure 3G**). The spikes observed in neurons indicate that neurons have developed functional receptors such as metabotropic receptors, ion pumps necessary for electrical signal transduction and synaptic transmission. Although spike frequency was similar between both conditions, neurons in CN+As nodes exhibited significantly narrower full-width half-maximum (FWHM) values for calcium peaks, suggesting shorter action potential durations and faster Ca^2+^ ion buffering (**Figure 3H**). This finding indicates that astrocytes contribute to neuronal functionality, potentially by enhancing calcium buffering capacity or accelerating functional neuronal maturation.

Taken together, these results demonstrate that integrating astrocytes significantly enhances node construction by improving network organization within the node, stabilizing node shape, and supporting neuronal functionality.

### Neuronal node Formation in Ex Vivo Mouse Cortical Cells

To validate the optimized parameters and procedures for neuronal node formation established in the human iPSC-derived model and assess the platform’s applicability to mouse *ex vivo* model, which fully developed neuronal properties, we created nodes using either human iPSC-derived cortical neurons and astrocytes or dissociated cells from mouse cortices. Unlike iPSC-derived cells, *ex vivo* mouse cortical cells contain not only neurons but also a mixture of glial cells and ECM components. Based on the human iPSC-derived node, we estimated the seeding cell numbers equivalent to 8x and 16x packing densities for mouse neuronal node formation. As expected, human neuronal nodes with 16x packing were successfully formed (**Figure 4A**). Interestingly, dissociated mouse cortical cells also established nodes that followed the plating device geometry. These nodes appeared to consist of multiple clusters averaging 300 μm in size (+/-90 μm SD), arranged in a linear configuration. Notably, mouse neuronal nodes seeded at 8x condition produced a similar node size to human CN+As nodes at 16x packing (**Figure 4B**). However, at 16x packing, mouse nodes displayed less consistency, with cells accumulating more densely at the center of the node (**Figure 4C**). This suggests that fewer cells are required for mouse node formation compared to human iPSC-derived nodes, potentially due to the endogenous ECM present in ex vivo cell dissociation, which may enhance cell cohesion. Additionally, the two node types exhibited slightly different characteristics. While MAP2-positive neurons from human iPSC-derived nodes showed some migration around the node, mouse cortical neurons predominantly remained better aggregated within the node. Conversely, astrocytes and other unidentified cells from mouse cortices migrated to distance of up to 600 μm from the node (**Figure 4D**). Altogether, these findings demonstrate that ex vivo mouse cortical cells can successfully form nodes using our bioengineered platform, showing similarities to human iPSC-derived neurons while exhibiting unique behaviors due to their native cellular and ECM composition.

**Figure 4.**
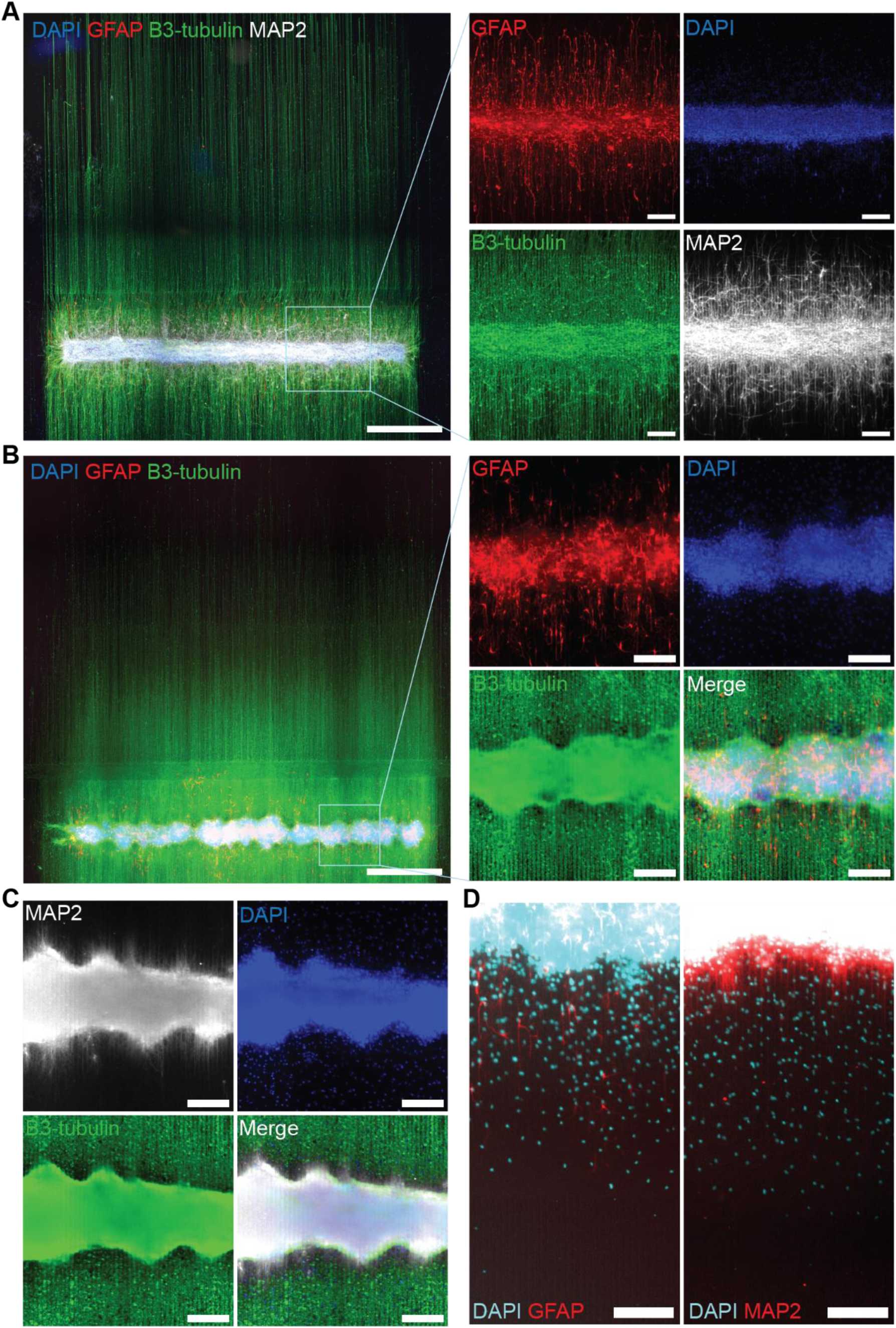
Validation of neuronal node construction in mouse ex-vivo culture. A) Neuronal node derived from human iPSC cortical neurons and astrocytes plated at 16x hexagonal packing, scale bar: 1000 μm. The dashed square shows a zoom-in view showing the organization of astrocytes (GFAP), nuclei (DAPI), axons (β3-tubulin) and dendrites (MAP2) within the node, scale bar: 200 μm. B-D) Neuronal nodes from mouse ex vivo neurons E16.5 plated at different hexagonal packing: B) Node plated using cell numbers calculated for 8x hexagonal packing, scale bar: 1000 μm. The dashed square shows a zoom-in view showing the organization of astrocytes (GFAP), nuclei (DAPI), and axons (β3-tubulin), scale bar: 200 μm. C) Node plated using cell numbers calculated for 16x hexagonal packing, stained for dendrites (MAP2), nuclei (DAPI) and axons (β3-tubulin). Scale bar: 200 μm. D) Mouse ex vivo neuronal nodes showing MAP2-positive neurons predominantly contained within the node, with some astrocytes and unidentified cells migrating around the periphery. Scale bar: 200 μm.

### T-Junction Microtopography Facilitates Neurite Diversion for Unidirectional Circuit Assembly

The randomness inherent in conventional *in vitro* neuronal network cultures can hinder accurate modelling of *in vivo* circuit development, as neuronal connectivity in the brain is highly organised and specifically connected from one area to another. Achieving precise network directionality *in vitro* significantly expands experimental capabilities for studying various aspects of synapse formation and function. To address this, we developed a microtopographical design featuring T-junction patterns to guide neurite directionality and facilitate unidirectional circuit construction.

The T-junction microtopography consisted of perpendicular microgroove arrays (10 µm height × 5 µm width) intersecting vertical grooves every 3 mm to accommodate multiple node assemblies (**Figure 5A**). Previous studies have shown that axons encountering such features can either turn to follow the cue or cross over to continue their original directions^17^. To identify the minimal T-junction width required to effectively divert axons without excessive spatial separation of overlapping axons from neighboring nodes, we tested three T-junction designs: 250 µm grooves (T1), 500 µm grooves (T2), and a combination of 250 µm grooves and 250 µm flat surfaces (T3). These designs were compared to a control containing only stripes of flat surfaces between vertical grooves (**Figure 5B**). Human iPSC-derived cortical neurons and astrocytes were plated beneath the T-junction areas, and neurite outgrowth was monitored using live-cell imaging with Sir-tubulin staining. After one week, in the control condition, most neurites extended directly across the flat surface without significant diversion (**Figure 5C**). Conversely, in the T-junction designs, neurites were concentrated at the T-junction, with some turning perpendicularly and others extending into neighboring grooves. Among the three designs, T1 exhibited the highest number of axons crossing the T-junction, whereas T2 and T3 better confined neurites within the T-junction region (**Figure 5D**).

**Figure 5.**
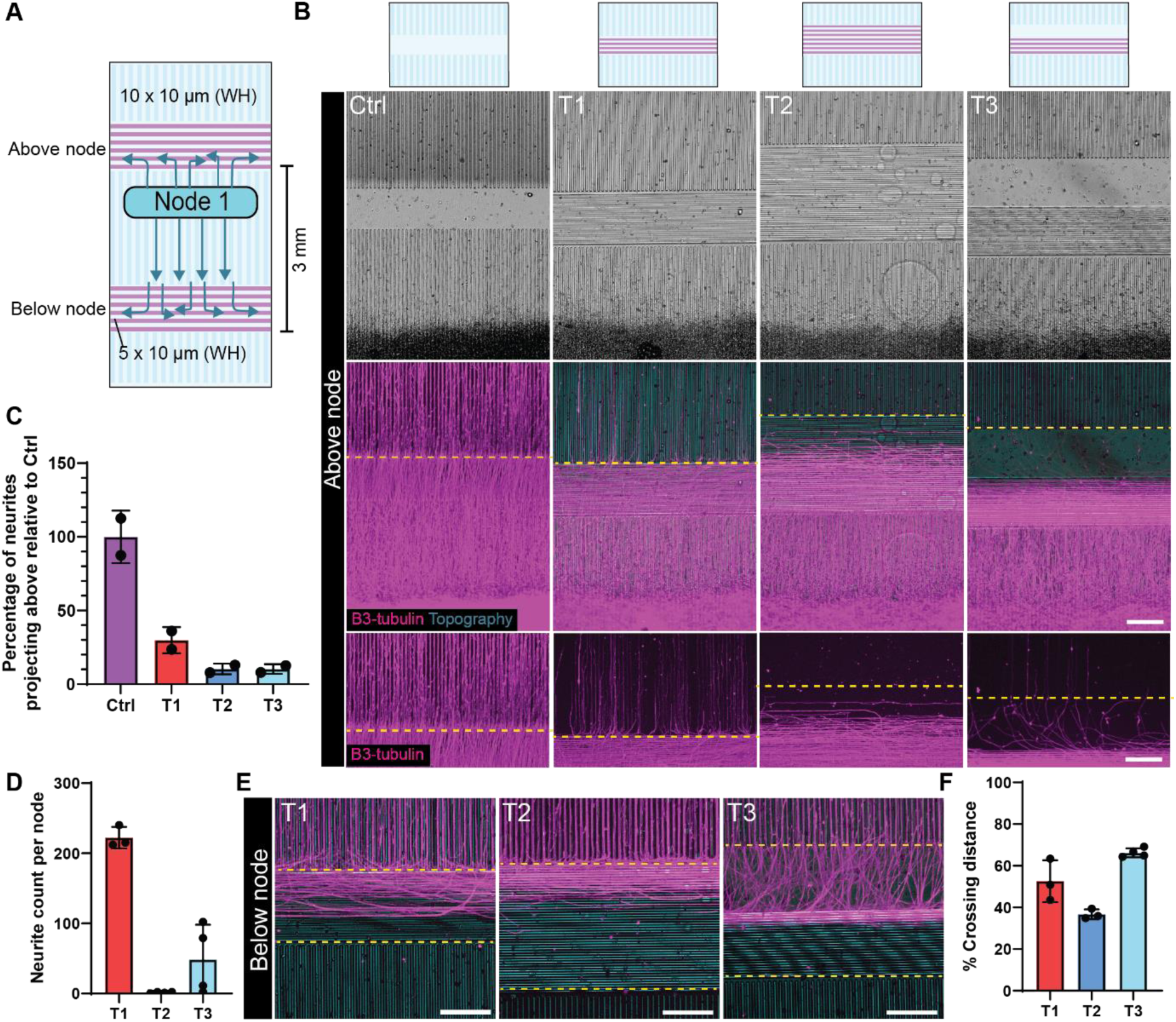
Microgroove topography directs neurites to the sides, allowing for unidirectional circuit construction. A) Schematic diagram of T-shape micropatterns to control neurite direction for unidirectional circuit. B) Representative images of T-shape design with neuronal node plated underneath the horizontal grooves (T-junction: T1, T2, T3) and control (Ctrl) without horizontal grooves. Neurites were stained with β3-tubulin at 7 days after plating. Scale bar: 200 μm. C) Percentage of neurites projecting upward based on integrated intensity of β3-tubulin in areas above T-junction (yellow dashed line). Data points represent the average per well, each containing 1-3 nodes. D) The number of neurites projecting above the T-junction (yellow dashed line). Data points represent the measurement from an individual node E) Representative images of neurites focusing on T-junction below the node area. Scale bar: 200 μm. F) Percentage of distance coverage of axons on T-junction below the node area. Data points represent the average of 6 measurements per node. All analyses in this figure were conducted on 1 independent experimental block, with 2 technical well replicates, each containing 1-3 nodes.

At the T-junction below the nodes - where neurites from two nodes would meet and connect - perpendicular grooves diverted axons extending from vertical grooves in all conditions (**Figure 5E**). The inclusion of flat surfaces in T3 further facilitated neurite turning before entering the grooves, allowing neurites to span a broader area (**Figure 5E, F**). This would enhance overlap between axons from two nodes in a pre-to-post-node layout. Considering both factors—minimizing axonal crossover and promoting neurite overlap—we selected T3 as the optimal pattern for constructing unidirectional circuits on our platform.

### Construction of a Large-Scale Cortical Circuit with Precisely Controlled Node Positions and Connectivity Sites

To construct a cortical circuit with controlled node positions and connectivity sites, we utilized the unidirectional layout described previously. This design enabled the assembly of two neuronal nodes aligned with the T-junction microtopography (**Figure 6A**). To achieve this, we developed a plating device consisting of two parallel wells, each with 247.5 µm × 4,950 µm (width x length), spatially aligned with the distance between two T-junctions (**Figure 6B**). The plating device mould was generated using SLA printing and subsequently used to produce the plating device. The T-shape micropattern was fabricated on a silicon master via photolithography and used to create PDMS micropatterned substrates, which were then biofunctionalised by O^2^ plasma treatment followed by PDL and laminin coating (**Figure 6C**). Before cell seeding, the surface of the plating device was activated using O^2^ plasma to enhance cell entry into the wells. The plating device was then assembled onto a biofunctionalized PDMS topographical substrate, precisely positioning each well beneath a T-junction (**Figure 6D**). Cortical neuron progenitors were co-seeded with astrocyte progenitors into each well. To prevent cross-contamination between the two nodes, the liquid from each well was carefully isolated during the plating process. The cells were confined within the plating device for 48 hours to allow aggregate formation, after which the device was removed. This resulted in a barrier-free platform for cortical circuit formation, allowing the differentiation of neurons and the extension of neurites guided by the underlying topography to establish network connectivity (**Figure 6D, E**).

**Figure 6.**
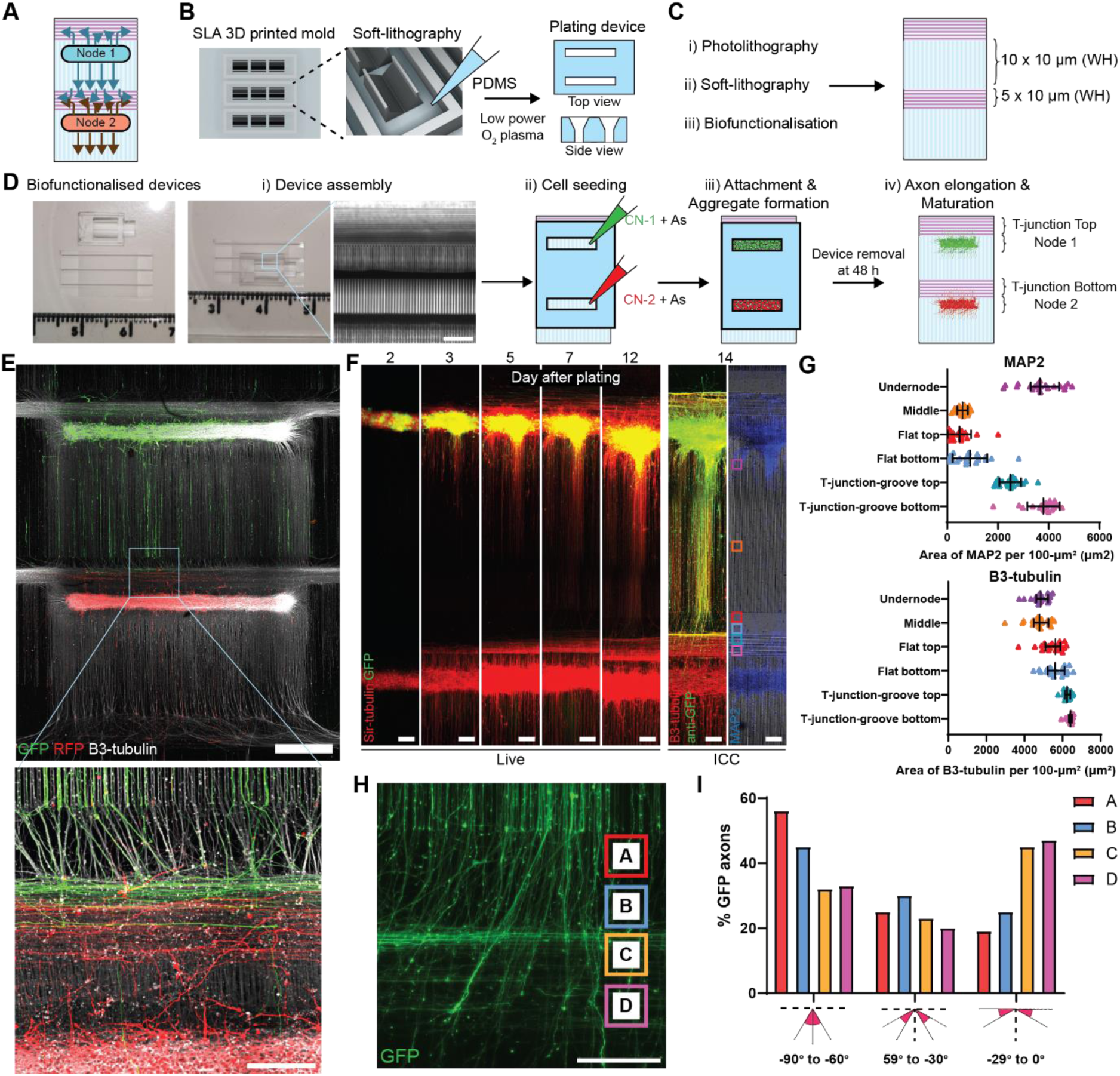
Assembly of cortical circuits with two nodes connecting their neurites at the T-junction. A) Schematics of BIOCONNET platform showing topographical cues that control neurite directionality to bias overlapped neurites from two nodes at T-junction. B) PDMS stencil of 2-pocket plating device is generated from 3D-printed mould and exposed to O^2^ plasma before cell plating. C) PDMS stencil of T-shape micropattern is generated from photolithography-based mould and biofunctionalized with O^2^ plasma, coated with PDL and laminin before cell plating. D) The plating device is placed on the micropattern, in which the pocket aligned approximately 200-250 µm below horizontal grooves (i). Scale bar: 200 µm. For cell seeding, two cortical neuron progenitor populations, mixed with astrocyte progenitors are seeded into each pocket and allowing aggregate formation for 48 hours before the device removal. Cells undergo neurite extension and network formation under differentiation and maturation processes. E) Overview of cortical neuron circuits between GFP and RFP cortical neuron node. Scale bar (top): 1000 µm. A zoom-in at the T-junction shows overlapping neurites from both nodes. Scale bar (bottom): 200 µm. F) Representative images of GFP to non-GFP cortical circuits. Neurite tracing labeled by Sir-tubulin shows the extension of axons from the node, imaged after device removal at day 2 post plating. After 14 days of differentiation, cells were fixed and stained with anti-GFP, MAP2 and β3-tubulin, showing the spatial arrangement of dendrites and axons labelled by MAP2 and β3-tubulin, respectively. Scale bar: 200 µm. G) Quantification of the area of dendrites (top) and axons (bottom) per 100-μm^2^ at different locations along the circuit, as indicated by representative squares in F. Data points represent ROIs cropped across circuits, expressed as median ± interquartile range, from 1 independent experimental block with 1 cell line each with 2 technical replicates. H) Representative image of T-junction area between two nodes, showing the alignment of GFP cortical neurons’ axons from the top node while approaching the neurites from the bottom node at 14 days of differentiation. Scale bar 200 µm. I) Directionality of GFP-positive axons across different locations indicated in H. The directions aligned vertically are defined as 90°, while horizontally defined as 0°. Data point represents the average measurement from ROIs cropped across 2 circuits, from 1 independent experimental block with 1 cell line each with 2 technical replicates.

This system is highly flexible, allowing the construction of circuits with distinct neuronal populations in each node. For instance, cortical neurons with different fluorescent markers, such as GFP-expressing neurons in one node and RFP neurons in the other, can be used to distinguish neurite outgrowth visually (**Figure 6E**). Alternatively, entirely different neuronal subtypes could be seeded in each node to customize the circuit’s functionality.

Additionally, the platform is compatible with live imaging, enabling real-time tracing of neurite dynamics during cortical circuit formation. To observe neurite outgrowth and connectivity, we constructed circuits using GFP-expressing neurons in one node and non-GFP neurons in the other, combined with Sir-tubulin live staining to trace neurite projections (**Figure 6F**). Over 6-7 days in culture, neurites extended from the nodes toward the T-junction area, following the microgroove topography. At the T-junction, neurites from both nodes began to accumulate and overlap, creating potential connectivity sites where synaptic connections could form (**Figure 6E, F**).

We further analyze the spatial distribution of dendrites and overall neurites within the circuit using MAP2 and β3-tubulin staining. MAP2-positive dendrites were predominantly concentrated around the nodes and within the grooves of the T-junction near the lower node, with their presence gradually decreasing further away from the node (**Figure 6G**). In contrast, overall neurites showed a slight increase at the T-junction area, likely due to the accumulation of neurites from both nodes. This spatial organization indicated that at the T-junction, axons from the top node likely overlapped with both dendrites and axons from the bottom node, reinforcing the potential for synaptic connectivity in this area. This platform demonstrates precise spatial arrangement for circuit construction.

To examine how axons from the top node interact with and span the area at the T-junction in the presence of neurites from the bottom node, we analyzed the directionality of GFP-expressing axons as they entered the T-junction. Approximately 45% of axons turned to follow the horizontal grooves of the T-junction, 30% continued vertically against the topographical cues, and the remaining axons exhibited intermediate directions (**Figure 6H, I**). This observation suggests that axons from the top node not only align with the horizontal grooves but also maintain vertical growth, effectively spanning the T-junction area, therefore enhancing higher chance for synaptic connectivity between nodes.

### Synaptic Connectivity Between Nodes on the Platform

Apart from the overall circuit visualization, the platform allows high-resolution observation at synaptic level in a large scale, both in live and fixed samples. This allows us to observe axonal dynamics during synaptic formation and quantify synaptic puncta after fixation and immunostaining. Additionally, the platform supports proteomic profiling, allowing for the identification and analysis of proteins associated with synaptic connectivity.

To observe the dynamics of synaptic connectivity between the two nodes, we conducted live imaging focused on the T-junction, where overlapping neurites from both nodes were previously observed. Imaging was performed when neurites first reached the T-junction and after 3 weeks of maturation (**Figure 7A, B**). Cortical circuits were constructed using GFP- and RFP-expressing cortical neurons, allowing us to distinguish neurites originating from each node. On day 6, as axons from the top node reached the T-junction, live imaging was conducted every 30 minutes over four hours. Axonal growth cones displayed dynamic pathfinding behaviors, some axons paused upon encountering RFP-positive neurites, forming bouton-like structures, possibly indicative of initial synapse connections. Others transiently formed such structures before continuing to navigate for other targets. By 3 weeks post-plating, the circuits exhibited greater overlap between GFP and RFP neurites, with more refined and interconnected structures, indicating network establishment. Interestingly, even in this mature state, axonal growth cones displayed similar dynamic behaviors, such as pausing and interacting with neighboring neurites. These observations demonstrate the platform’s ability to capture real-time axonal dynamics with single-axon resolution during cortical network formation and in more matured cultures, making it a promising platform for studying axonal interactions and connectivity between defined neuronal populations.

**Figure 7.**
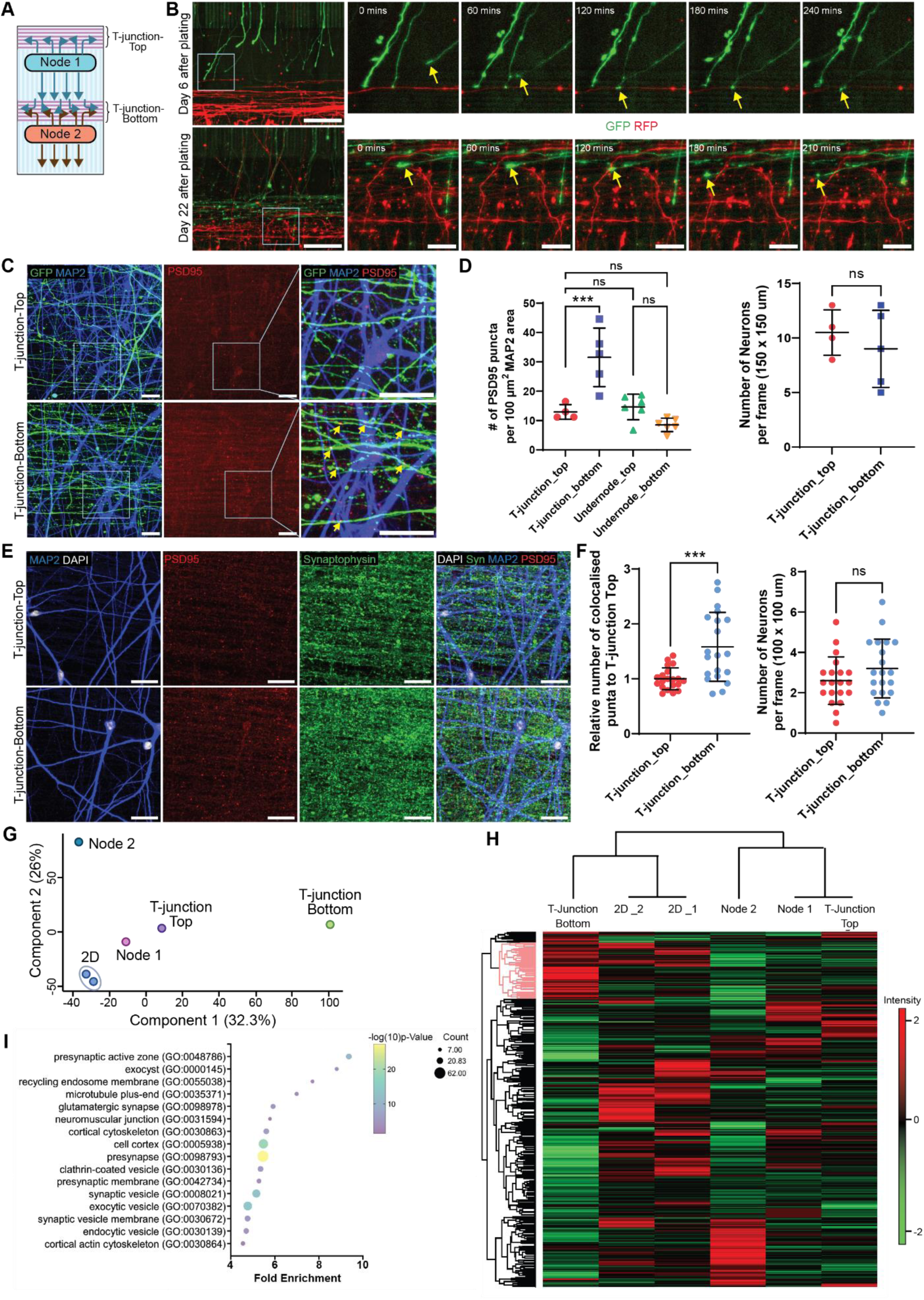
Assessment of synaptic connectivity between two nodes. A) Schematics of BIOCONNET platform. B) Representative images of circuit formation between axons of GFP-cortical neurons from the top node and neurites of RFP-cortical neurons from the bottom node observed via live-imaging at day 6 and day 22 of differentiation. Scale bar 200 μm for the overview and 50 μm for the series of zoom-in views. C) Representative images at the T-junction-top and T-junction-bottom area of the circuits with GFP neurons at the top node connecting with no-fluorescence neurons at the bottom node. An increase in PSD95 expression is observed at the T-junction-bottom with PSD95 puncta spotted on dendrites overlapping with GFP-axons from the top node (yellow arrow). Scale bar: 20 μm for the overview and 10 μm for zoom-in views. D) Quantification of the number of PSD95 puncta per 100 μm^2^ MAP2 positive dendrite areas (left) and the number of neurons per frame of analysis (right). Data points represent ROIs imaged spatially across a circuit from 1 independent experimental block with 1 cell line, one-way ANOVA with Tukey’s multiple comparisons, mean ± SD, ***P < 0.001, ns: non-significant. E) Representative images at the T-junction-top and T-junction-bottom area, showing synaptic protein expression PSD95 and synaptophysin, with dendrites and nuclei staining used for the analysis. Scale bar: 20 μm. F) Quantification of the number of colocalised puncta per 100 μm^2^ MAP2 positive dendrite area. Data points represent FOVs imaged spatially across 5 circuits, from 3 independent experimental block with 2 cell lines, each with 1-2 technical replicates, unpaired t-test to test for significance, mean ± SD, ***P < 0.001, ns: non-significant. G) PCA of isolated fractions, showing clear separation between T-junction Bottom (by Component 1) and Node 2 (by Component 2). H) Hierarchical clustering of samples using Pearson correlation and proteins using Maximum distance. The cluster identifying upregulated proteins is highlighted in red. I) Top 16 GO terms with the highest fold enrichment for the selected cluster (highlighted in red in H) using overrepresentation analysis in PANTHER.

To confirm the sites of synaptic connectivity between nodes, we performed spatial quantification of post-synaptic density 95 (PSD59) puncta along the circuits. For this, GFP cortical neurons and non-GFP cortical neurons were used to construct the circuit. After allowing the circuit maturation for 28 days, cultures were fixed and stained for PSD95, GFP, and the dendritic marker MAP2. We observed higher PSD95 expression at the T-junction of the bottom node compared to the top node. Notably, many PSD95 puncta were found to overlap with MAP2-positive dendrites and GFP axons, suggesting synaptic connectivity between the two nodes (**Figure 7C**, **Supplement Figure 3C**). The puncta within the dendritic area were quantified using a customized semi-automated pipeline (**Supplement Figure 3A**). The result showed that the number of PSD95 puncta per 100 µm^2^ of MAP2-positive dendritic area was approximately 2-fold increased at the T-junction of the bottom node compared to the T-junction at the top node (**Figure 7D**). No significant differences were observed in PSD95 among the T-junction of the top node and both under node positions, which served as internal controls as indicative of the intra-node synaptic basal levels. In addition, there was no significant number of neurons occupied at both the T-junction top and bottom, suggesting unbiased confounding effect from neuron numbers. Altogether, this indicates that the increase in PSD95 puncta at the T-junction-bottom is influenced by connectivity between the two nodes, confirming inter-node synaptic formation at this position.

To further validate synaptic connectivity between two nodes at the T-junction, we quantified the colocalization of pre- and post-synaptic protein puncta, synaptophysin and PSD95. Cortical circuits were constructed with the same population of human iPSC-derived cortical neurons and astrocytes and differentiated for 21-28 days to ensure synaptic formation and stability of synapses (**Supplement Figure 3B**). The circuits were fixed and stained for synaptophysin, PSD95, MAP2 and DAPI. Colocalized puncta were quantified on MAP2-positive dendrites, expressed as the number of puncta per 100 µm^2^. The quantification revealed a significant increase in colocalized synaptophysin and PSD95 puncta at the T-junction of the bottom node, compared to the top node. Importantly, the number of neurons within the analyzed area was consistent between T-junctions, ruling out cell density as a contributing factor. (**Figure 7E, F**). These results indicate that the cortical circuits formed on the platform exhibit increased synaptic connectivity between nodes.

A major advantage of this large-scale, open system is its ability to isolate specific circuit regions for proteomic analysis. As a proof of principle, we performed proteomic profiling across the circuits to determine whether synapse-related proteins could be identified at predefined sites of synaptic connectivity. To achieve this, we dissected and extracted proteins from the circuit regions: T-junction-top, Node 1, T-junction-bottom, and Node 2 (**Supplementary Figure 4A**). Cortical neurons were cultured on bioengineered platforms for 28 days before protein extraction, with conventional 2D cultures processed in parallel as controls. Protein yields from each region (pooled from 10 circuits) or from three wells of a 12-well plate for 2D culture were sufficient (∼10 µg) for mass spectrometry (MS) processing and analysis. Quantification of proteins identified from each isolated region, with ∼200 ng loaded per sample, revealed comparable protein counts (∼8,000–10,000 per region; **Supplementary Figure 4B**). To identify similarities and differences across the circuits, we applied principal component analysis (PCA). The analysis revealed that the T-junction-bottom, where inter-node synapses form, was distinctly separated from other regions along principal component 1. In contrast, Node 1, the T-junction-top, and 2D cultures clustered closely together, while Node 2 was segregated along principal component 2 (**Figure 7G, Supplementary Figure 4C, 4D**). Additionally, hierarchical clustering analysis showed that the protein expression profile of the T-junction-bottom was distinct from other regions. Especially upregulated protein clusters (in red) in the T-junction bottom, show lower expression in Node 2, Node 1, and the T-junction-top (**Figure 7H, Supplementary Figure 4E**). Gene ontology (GO) analysis of this protein cluster upregulated at the T-junction-bottom revealed a strong association with synaptic components, including the presynaptic active zone, glutamatergic synapses, and synaptic vesicles (**Figure 7I**). These findings demonstrate that the platform enables proteomic analysis and confirms synaptic connectivity between nodes. By allowing precise isolation of circuit compartments and subsequent proteomic analysis, this system serves as a powerful tool for investigating the molecular composition of synapses within defined neuronal networks.

### Modulation of Human iPSC-Derived Cortical Network Connectivity Using Optogenetic Tools

To assess whether the engineered circuits exhibit functional connectivity, we employed optogenetic tools in combination with calcium imaging. Cortical circuits were established with ChrimsonR-expressing cortical neurons – a red-light drivable channelrhodopsin - at the top node and wild-type cortical neurons at the bottom node (**Figure 8A**). ChrimsonR-expressing neurons can be stimulated with red light, triggering action potentials that propagate synaptic signals to depolarize connected neurons. This neuronal activity was monitored using Fluo-4AM, a calcium sensitive dye, allowing simultaneous modulation and observation of neuronal activity. Cortical neurons were terminally differentiated for 21 days before imaging experiments. Baseline neuronal activity was first recorded via calcium imaging, followed by optogenetic stimulation using a 100 ms LED pulse at 585 nm delivered every 30 seconds. Due to limitations in dye penetration and light path within the 3D structure of the nodes, the analysis focused on neurons located near the nodes as a representative population. Calcium peaks in neuronal somas were detected and presented as trains of spikes using PeakCaller software^18^. The results revealed that neurons at both nodes exhibited spontaneous neuronal activity (**Figure 8B**, i, iii). Following optogenetic stimulation, the same neurons were analyzed and showed that neurons in the top node changed from random spike trains to stimulus-driven peaks, corresponding to the stimulation pulses (**Figure 8B**, ii). However, neurons in the bottom node exhibited more diverse changes in spike patterns. In particular, peaks in the wild-type node occurred with a delay relative to the stimulation time points (**Figure 8B**, iv), suggesting that neuronal firing patterns in the bottom node were modulated through synaptic connectivity with the ChrimsonR-expressing neurons in the top node. These results indicate that cortical circuits on the platform may establish functional connectivity, with optogenetic stimulation providing precise control over neuronal activity. This demonstrates the platform’s potential for investigating activity-dependent processes within cortical networks.

**Figure 8.**
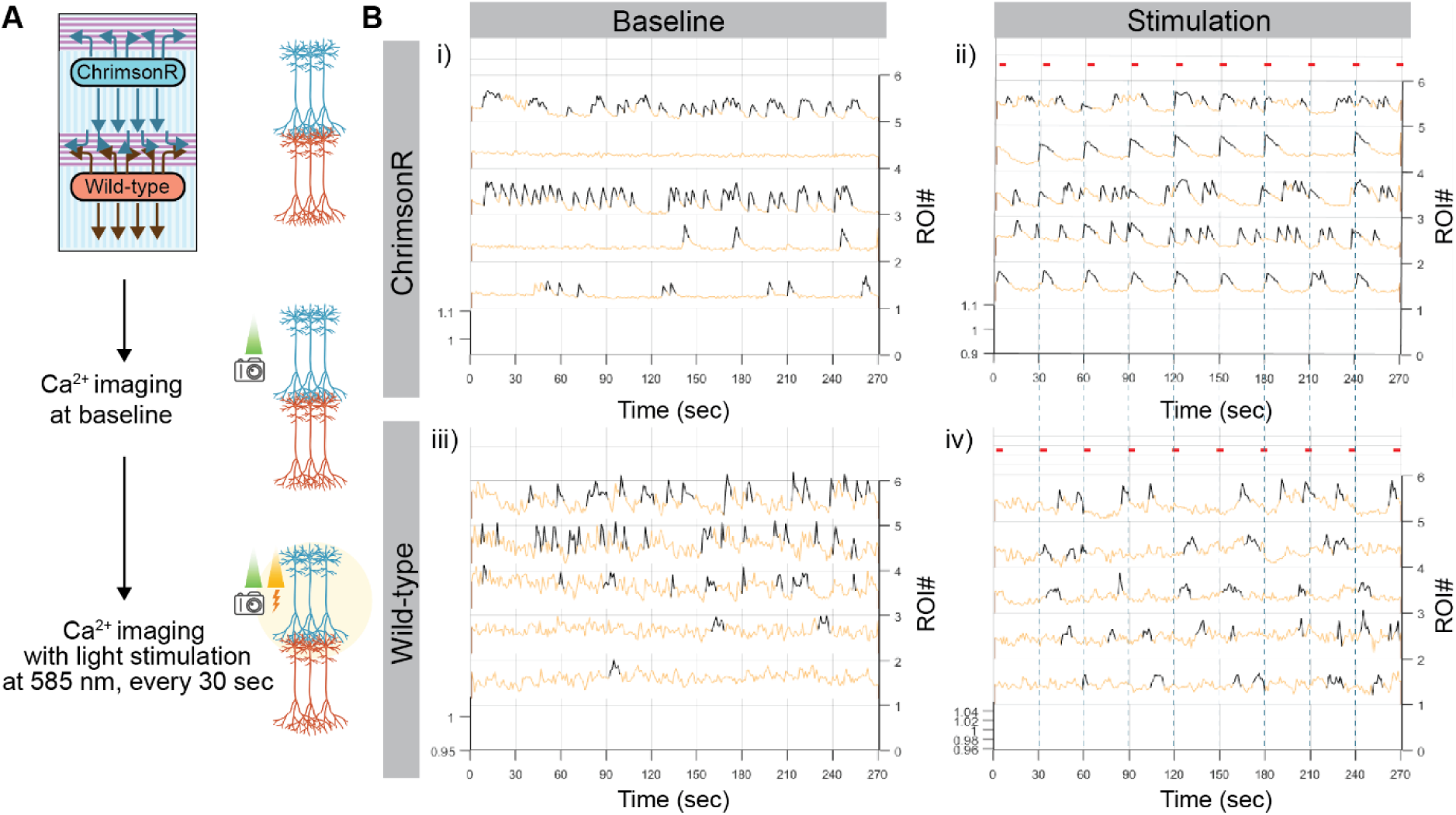
Integration of optogenetic expressing cortical neurons allows for network activity alteration. A) Schematics illustrating experimental processes, where cortical circuits connect a node expressing the optogenetic tool ChrimsonR to a wild-type node. Baseline of calcium signals were acquired through time-lapse imaging of calcium dye Fluo-4AM (490 nm), followed by calcium imaging with optogenetic stimulation (585 nm, 100ms pulse) every 30 seconds. B) Calcium traces of each ROIs from the ChrimsonR node (i, ii) and the wild-type node (iii, iv), with the time points at which stimulation was triggered shown in dashed lines and red dashes.

## 4. Discussion

*In vitro* neuronal cultures provide simplified platforms for studying neurobiology and modeling disease. However, their inherent lack of complexity, particularly the absence of defined network architecture, imposes several limitations^4^. This lack in organization at the multicellular level restricts investigations into intricacies of network formation, activity-dependent processes^19^ and neural network functions.

To address these challenges, we developed a cortical circuit platform that integrates bioengineering techniques with iPSC-derived cortical neurons. This platform provides the ability to systematically control and alter the patterns of connectivity. By employing temporary cell confinement with optimal cell numbers, cortical neurons self-organize into spatially defined nodes on a topographic micropattern. This design allows for the construction of defined circuit architectures without physical barriers, creating an open platform that supports precise spatial organization of circuits. Using this system, we showed that the platform enables flexible adjustments in node geometry and cellular composition, facilitating the modeling of diverse circuit complexities.

Our approach combines the strength of 2D (simplicity and accessibility) and 3D cultures (more physiological relevance). Unlike 3D organoids or *in vivo* models, which have limited control over intrinsic connectivity, our platform employs 3D neuronal nodes guided by 2D substrate topography to direct axonal paths and synaptic connectivity. This hybrid structure supports systematic analysis of synapse formation at predetermined sites, advancing our ability to investigate the mechanisms underlying circuit formation.

Additionally, the platform is compatible with live-imaging systems, allowing versatile imaging at various magnifications from large-scale circuit overviews to high-resolution imaging via wide-field and confocal microscopy. This supports diverse applications of molecular and cellular studies, including tracking axon growth, monitoring synapse formation, and observing neuronal calcium dynamics. When combined with optogenetics, the platform also facilitates controlled modulation of neural activity, making it a valuable tool for cortical network studies.

Importantly, the open design of our platform overcomes key limitations of microfluidic devices, which are constrained by their enclosed systems. Microfluidics often lack scalability and accessibility, limiting their utility for experiments requiring substantial cellular material, such as transcriptomics or proteomics. In contrast, our open system accommodates larger cell quantities and circuit scales, thereby enabling spatial isolation of specific circuit compartments for RNA or protein extraction. This capability opens new opportunities for investigating molecular and cellular events within specific circuit components at defined developmental stages. For example, the platform could be used to study molecular events changes at axon terminals during growth and synapse formation by combining with omics analyses.

The platform is also well-suited for human disease modeling, addressing the need for human-specific models to investigate molecular and cellular mechanisms that may differ across species^20^. Its ability to manipulate synaptic connections between diseased and healthy neurons allows for investigations into synaptic loss and network dysfunction in conditions such as neurodegeneration. For example, alterations in connectivity between neuronal nodes could be used to examine how synaptic loss contributes to network dysfunction and cell death, or to investigate the underlying cellular mechanisms-whether connectivity defects arise from impaired cellular processes such as axonal transport, or whether defective cellular functions drive synaptic loss.

Despite its advantages, the platform presents some limitations compared to microfluidic systems. The absence of physical barriers exposes all neuronal populations to the same media, making it unsuitable for experiments that require different media or media gradients. In addition, the open system presents the issue of potential cell migration, as observed in this study, where cortical neurons were found surrounding the node. This migration may restrict experiments requiring precise isolation of somas and axons or pure inter-node synaptic elements without contamination of neuronal soma from one node. Furthermore, the system requires a threshold number of cells to form compact and homogeneous neuronal nodes, which limits its scalability for small-scale circuits compared to microfluidic devices.

Although these challenges pose some constraints, specific strategies could mitigate the platform’s limitations. The large-scale design of the platform is particularly advantageous for substrate manipulation, by allowing selective coating of specific areas with molecules for targeted purposes. For instance, pre-treatment of T-junction areas with semaphorin 3A, which promotes dendrite over axon development^21^, could potentially enhance compartmentalization within circuits. In addition, the migration issue could be mitigated by pre-treating neurons under differentiation conditions to promote their transition to the post-mitotic stage before seeding. This approach would initiate the maturation process earlier, reducing migration, a characteristic of immature early post-mitotic neurons^22,23^ . Similarly, ex vivo mouse neuronal nodes, which develop faster maturation process than human neurons, exhibit better coherence and minimal migration.

Finally, the platform offers a promising foundation for studying cortical circuit formation and function, with opportunities to refine and expand its applications. Future improvements could focus on integrating additional biological complexities, such as incorporating inhibitory neurons, which are essential for excitatory/inhibitory balance, cortical circuit function and maturation^24^. Moreover, the platform’s flexibility in manipulating circuit architecture could be extended to increase network complexity by adding more nodes and adjusting circuit configurations. This would enable investigations into the hierarchical organization of modular systems^25,26^ to gain insight into cortical network function.

## 5. Conclusion

Overall, our cortical circuit platform presents significant advancements over traditional 2D *in vitro* models and microfluidic technologies. This study demonstrates the platform’s capability to construct scalable circuits without the need for physical barriers. This platform enables the reproducible creation and modification of circuit architectures, offering control over spatial organization, the ability to track axonal growth, and systematic observation and analysis of synapse formation. Furthermore, its scalability supports the retrieval of circuit materials for molecular analysis, including proteomic analysis, providing new opportunities to investigate the processes underlying circuit formation and synapse loss in health and disease conditions.

## 6. Experimental Section

### Cell Culture

All cells were cultured in a humidified incubator at 37°C with 5% CO^2^.

#### Human iPSCs culture

Human iPSCs were cultured on plates coated with Matrigel (Corning, 1:100) in Essential 8 Flex medium (Gibco) and passaged using EDTA-PBS (Gibco, 0.5 mM). Details of the iPSC lines are provided in **Table 1**.

**Table 1.** Human iPSC lines.

| Cell line name | Donor Age | Donor Sex |
| --- | --- | --- |
| Control 3 | Unknown | Female |
| KOLF2C1, KOLF2.1J | 55-59 years old | Male |

#### Generation of cortical neuron progenitors

Cortical neuron progenitors were generated using dual SMAD inhibition following a previously published protocol^12^ with some modifications (**Supplementary Figure 1**). Briefly, iPSCs were cultured to 100% confluence before induction. Cells were fed daily for 8-10 days with a neural induction medium containing SB431542 (Tocris, 10 µM) and Dorsomorphin (Merck, 1 µM) in a base medium composed of 50% advanced Dulbecco’s modified eagle’s medium (DMEM) (Gibco) and 50% NeuroBasal (Gibco) with B27 (Gibco) and N2 (Gibco) supplements, Penicillin-Streptomycin antibiotic mix (Gibco, 100 μg/ml) and GlutaMAX (Gibco, 1x).

At this stage, iPSCs were transitioned to neuroepithelial cells. They were passaged using EDTA-PBS (Gibco, 0.5 mM), centrifuged at 100 x g, and reseeded 1:2 on laminin-coated plates (Sigma, L2020, 10 μg/ml) in base medium. To promote rosette formation, Matrigel (Corning, 1:5) was applied on the cell surface for 30 minutes, allowing the Matrigel to gel into a thick layer before topping up with culture medium. After 2-3 days, upon appearance of neural rosettes, cells were passaged on laminin-coated plates with a Matrigel layer, in base medium supplemented with FGF (Gibco, 20 ng/ml) (F20 medium).

This process continued until passage 3, where cortical neuron progenitors arranged in rosette structures were enriched. After passage 3, progenitors were passaged using EDTA-PBS, centrifuged at 400 x g, and plated on Matrigel-coated plates (1:100) without additional Matrigel layer. Cortical neuron progenitors were cryopreserved in 10% DMSO in F20 medium. For experiments using bioengineered platforms, cortical neuron progenitors from passage 7-24 (Day 21-95) were used. Specifically for synapse analysis, cortical neuron progenitors from passages 13-19 (Day 64-80) were used.

#### Neuronal differentiation

For terminal differentiation of cortical neuron progenitors, base medium supplemented with compound E (Enzo, 0.1 µM) was used. For conventional 2D culture, cortical neuron progenitors were plated on poly-D-lysine coated plates followed by laminin coating and maintained in the differentiation medium. A Matrigel spike (Corning, 1:100) was added the following day and every 7-10 days to support long term attachment.

#### Astrocyte progenitor culture and differentiation

Astrocyte progenitors were generated from human iPSCs according to a previously published protocol^27^. Cells were cultured on Matrigel-coated plates (1:100) in base medium supplemented with EGF (Gibco, 20 ng/ml) and FGF (Gibco, 20 ng/ml) (EF20 medium) and passaged using Accutase (StemPro). For differentiation, astrocyte progenitors were co-cultured with cortical neurons in base medium supplemented with compound E (0.1 µM).

#### Mouse ex vivo culture

Mouse ex vivo neurons were isolated from the cortices of E16.5 embryonic mice following a previously published protocol^28^. Dissociated neurons were cultured in bioengineered devices for node formation, initially in attachment medium containing Minimal Essential Medium (MEM) (Gibco), sodium pyruvate (Gibco, 1 mM), and glucose (33 mM / 0.6%) for 5 hours. The medium was then replaced with maintenance medium containing Neurobasal (Gibco), B27 supplement (Gibco, 2%), GlutaMax (Gibco, 1%), Glucose (33 mM / 0.6%), and Penicillin-Streptomycin antibiotic mix (Gibco, 1x), and the plating device were removed 3 days later. Cultures were fed weekly and maintained in the maintenance medium for 14 - 21 days.

#### Transfection and selection of cortical neuron progenitors and human iPSCs

Stable cortical neuron progenitor lines expressing GFP (PB-Ef1α-GFP-PGK-Puro) or RFP (PB-CAG-RFP-Hygro) were established for neurite visualization and circuit formation. A stable human iPSC line expressing ChrimsonR (PB-TetOn-hNGN2, EF1α-Puro-NLS-T2A-BFP, Syn-ChrimsonR-tdTomato) was created for optogenetic control.

Transfections were performed in Essential 8 medium (E8, Gibco) for human iPSCs and PenStrep-free F20 medium for cortical neuron progenitors. For both cell types, the desired *PiggyBac* (PB) construct and PB transposase were co-transfected to achieve stable genomic integration, using Translt-LT1 transfection reagent (Mirus). Cells were plated in 6-well plates one day before transfection. Each well received 2.5 μg of DNA in 250 µL of medium at a 1:3 ratio to the transfection reagent. The transfection mixture was incubated with cells for 24 hours before replacing it with fresh culture medium.

For puromycin selection, puromycin (Gibco) was initially used at 0.5 µg/mL for 48 hours post-transfection, then increased to 1 µg/mL. Several selection rounds were performed to achieve 100% BFP positive cells for iPSCs. For hygromycin selection, a concentration of 500 ug/ml hygromycin was used. After selection, cells were expanded, induced for cortical neuron differentiation, or used directly in experiments.

### Bioengineered circuit platform

#### Microfabrication of patterned substrates

Silicon masters with patterned substrates were fabricated using SU-8 photoresist via photolithography. The master with an array of 10 µm-wide grooves with 200 µm plateaus at every 5 mm were prepared using SU-8 2002 photoresist as previously described^11,29^. T-junction patterns composed of 10 µm-wide grooves (2.5 mm long) intersected perpendicularly with 5 µm-wide grooves, were prepared using SU-8 2007 photoresist.

For photolithography, silicon wafers were plasma-cleaned for 10 minutes and dehydrated by baking at 200°C for 2 minutes. Photoresist was spin-coated on the wafers at 1000 rpm for 40 seconds using a spin coater (Polos), creating a uniform layer. Soft baking was performed at 95°C for 2 minutes before photopatterning. Micropatterns were designed using CleWin software and written with a MicroWriter ML3 laser-based system at an exposure energy of 70 mJ/cm². Post-baking was carried out at 65°C and 95°C, followed by development with PGMEA to remove unexposed SU-8, and final cleaning with isopropyl alcohol (IPA). The new masters were silanised with Trichlorosilane (C^8^H^4^Cl^3^F^13^Si) in a vacuum chamber for 1 hour.

PDMS patterned substrates were fabricated using the Sylgard-184 silicone elastomer kit (Dow Corning). PDMS pre-polymer and curing agent were mixed at a 10:1 w/w ratio and degassed to remove bubbles. The mixture was spin-coated onto a patterned silicon master at 300 rpm with an acceleration of 1000 rpm/sec for 80 seconds using a Polos spin coater. The thin PDMS layer on the master was cured on a heat plate at 100 °C for 5– 10 minutes. The micropattern PDMS was then transferred to a clean plastic dish.

#### Fabrication of plating devices

Plating devices were designed using Autodesk Fusion 360 software as previously described^11^, with dimensions and sizes listed in **Table 2**. The designs were exported as.stl files and processed for slicing using Chitubox software with parameters optimized for 3D printing. The devices were fabricated using Phrozen Sonic mini 4K and Phrozen Sonic mini 8K 3D printers as previously described^11^.

**Table 2.** Plating device designs.

| Width (mm) | Length (mm) | Area (mm<sup>2</sup>) | Aspect Ratio (AR) | Used in Figure |
| --- | --- | --- | --- | --- |
| 0.0700 | 1.750 | 1.225 | 2.5 | 1F-G |
| 0.5000 | 2.450 | 1.225 | 4.9 | 2A |
| 0.4050 | 3.025 | 1.225 | 7.5 | 2A |
| 0.3500 | 3.500 | 1.225 | 10 | 2A, 2D-J, 3B-C |
| 0.2475 | 4.950 | 1.225 | 20 | 2J, 3D-H, 4, 5, 6, 7, 8 |
| 0.1750 | 7.000 | 1.225 | 40 | 2J |

### Biofunctionalisation

For micropatterned PDMS substrates, surfaces were oxygen plasma treated at 7 sccm with 100% power for 1 minute (Henniker Plasma). The treated PDMS was coated with poly-D-lysine (PDL) (Gibco) under UV light for 15 minutes, washed three times with PBS, and then coated with laminin (Sigma) at 37 °C for 2 hours. Laminin-coated PDMS substrates can be stored at 4 °C.

For plating devices cast from 3D-printed moulds, the top surfaces were oxygen plasma treated at 7 sccm with 50% power for 30 seconds prior to cell seeding.

#### Cell seeding

Biofunctionalised micropatterned substrates were placed inside 12-, or 6-well plates, washed with PBS and dried using an aspirator. Substrates were assembled with plating devices by placing the seeding well approximately 200-250 µm beneath the T-junction. The plating device was gently pressed onto the substrate with forceps to remove any trapped air. Cortical neuron progenitors and astrocyte progenitors were dissociated into single cells using Accutase (StemPro), centrifuged at 800 x g, and resuspended in 100 µl of differentiation medium with compound E (Enzo, 0.1 µM) per detached well to achieve a concentrated cell suspension. Cells numbers were counted and seeded per device as indicated in **Table 3**. To prevent drying, a few microliters of medium were added to the wells after seeding using a P20 pipette. Cells were allowed to settle for 2 hours in incubator before washing three times with PBS to remove unattached cells. Wells were then filled with differentiation medium. After 48 hours (unless stated otherwise) for aggregate formation, plating devices were removed using forceps, and cells were returned to the incubator for full attachment. A Matrigel spike (1:100) was added the following day and every 7-10 days to support long term attachment.

**Table 3.** The number of cells for seeding.

| Area (mm<sup>2</sup>) | Hexagonal packing | Cortical neurons (cells) | Astrocytes (cells) |
| --- | --- | --- | --- |
| 1.225 | 2x | 12,300 | 3,460 |
| 1.225 | 4x | 24,600 | 6,920 |
| 1.225 | 8x | 49,200 | 13,800 |
| 1.225 | 16x | 98,400 | 27,700 |

A hexagonal packing layer, denoted as x, was calculated based on the 2D arrangement of 15 µm circles, reflecting the average cell size, within a rectangular device, using the “Circles within a Rectangle – Calculator” available online^30^.

#### Immunocytochemistry, sample mounting, and live dyes

Cells were fixed with 4% PFA for 15 minutes, followed by three PBS washes.

For general staining (excluding synaptic and internal aggregate staining), cells were permeabilised with 0.1% TritonX in PBS for 10 minutes and blocked with 3% goat serum in PBS for 30 minutes at room temperature. Primary and secondary antibodies (details in **Table 4**, **Table 5**) were prepared in a solution of 0.05% TritonX and 1.5% goat serum in PBS. After blocking, cells were incubated with primary antibodies for 1 hour at room temperature, followed by three PBS washes. Secondary antibodies were used at a final concentration of 2 µg/ml and incubated for 30 minutes at room temperature. For nuclei staining, DAPI (Sigma) was included with the secondary antibody solution at a final concentration of 1 µg/ml.

**Table 4.** Primary antibodies.

| **Antibody** | **Host species** | **Isotype** | **Manufacturer** | **Catalogue code** | **Dilution** |
| --- | --- | --- | --- | --- | --- |
| TBR1 | Rabbit | Polyclonal | Abcam | Ab31940 | 1:300 |
| CTIP2 | Rat | IgG2a | Abcam | Ab18465 | 1:300 |
| BRN2 | Rabbit | Polyclonal | ThermoFisher | PA5-30124 | 1:500 |
| SATB2 | Mouse | IgG1 | Abcam | Ab51502 | 1:100 |
| β3-Tubulin | Mouse | IgG2b | Sigma | T5076 | 1:1000 |
| MAP2 | Mouse | IgG1 | Sigma | 13-1500 | 1:750 |
| OTX1/2 | Rabbit | Polyclonal | Abcam | Ab21990 | 1:200 |
| Nestin | Mouse | IgG1 | Merck | MAB5326 | 1:200 |
| PAX6 | Mouse | IgG1 | ThermoFisher | MA1-109 | 1:200 |
| GFAP | Chicken | Polyclonal | Abcam | Ab4674 | 1:2000 |
| GFAP-Cy3 | Mouse | IgG1 | Sigma | C9205 | 1:500 |
| Synaptophysin | Rabbit | Polyclonal | ThermoFisher | PA1-1043 | 1:500 |
| PSD95 | Mouse | IgG2a | ThermoFisher | MA1-045 | 1:500 |
| GFP | Rabbit | Polyclonal | Abcam | Ab6556 | 1:500 |

**Table 5.**
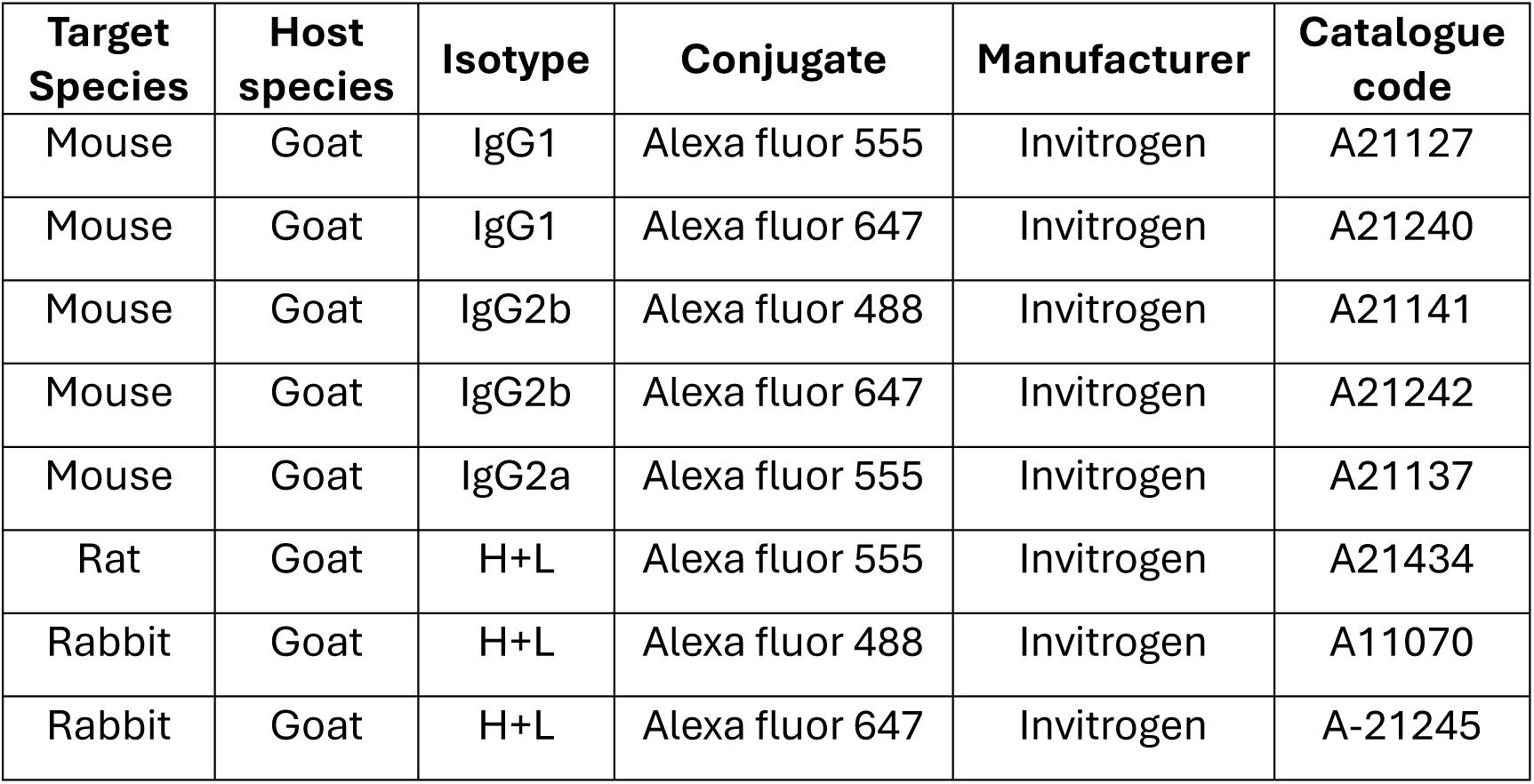
Secondary antibodies.

For synaptic staining, cells were permeabilized with 0.1% TritonX in PBS for 10 minutes and blocked with 3% goat serum in PBS for 2 hours. Primary antibodies were incubated overnight at 4°C, followed by two 20-minute PBS washes. Secondary antibodies were applied for 1 hour at room temperature, followed by two additional 20-minute PBS washes.

For internal aggregate staining, cells were fixed with 4% PFA for 1 hour, followed by three PBS washes. Permeabilized was performed with 0.1% TritonX in PBS for 2 hours, followed by blocking with 3% goat serum in PBS for 1 hour. Primary antibodies were incubated 24 hours at 4°C, followed by two 20-minute PBS washes. Secondary antibodies were incubated for 24 hours at 4°C, followed by two additional 20-minute PBS washes. DAPI diluted in PBS was incubated for 1 hour at room temperature, followed by two more 20-minute PBS washes. Samples were incubated overnight at room temperature in RapiClear 1.49 (Sunjin Lab) for optical clearing.

To mount samples on PDMS, PBS was removed, and the PDMS was transferred to an imaging slide using forceps. PDMS spacers were placed around the sample to prevent compression of 3D structures when maintaining the aggregate shape was required. FluorSave (Sigma) was applied on top of the sample, which was then covered with a coverslip #1.5.

Silicon rhodamine-tubulin (Sir-tubulin) (Spirochrome) was applied to cells at a concentration of 100 nM and incubated for 1 hour in the incubator, followed by a PBS wash to remove dye residues.

Fluo-4AM (ThermoFisher) was used at a concentration of 5 μM. Cells were incubated with the dye for 1 hour, washed with PBS, and replaced with fresh medium. Cells were rested in the incubator for at least 30 minutes before imaging.

### Experiments and image analysis

#### Characterisation of cortical neurons

To identify cortical neuron progenitors, cells were plated, fixed, and stained with neural progenitor markers Nestin and OTX1/2, along with DAPI. OTX1/2-positive cells were analyzed using a CellProfiler pipeline, which converted signals into objects and identified overlaps between OTX1/2 and DAPI. The percentage of OTX1/2-positive cells was calculated relative to the total cell count. Nestin expression was quantified by measuring total Nestin intensity in CellProfiler and normalizing it to the total cell count.

For cortical neuron subtype characterization, cells were terminally differentiated for 7 days, fixed, and stained with cortical neuron subtype transcription factor markers TBR1, CTIP2, BRN2, and SATB2. The percentage of nuclei expressing each subtype was analysed using CellProfiler software. DAPI and neuronal subtype channels were identified as objects and overlaps between DAPI and each subtype were detected. The percentage of each neuronal subtype was calculated relative to the total cell count.

To assess astrocyte presence, cells were terminally differentiated for 7 days, fixed, and stained with the GFAP marker. GFAP-positive cells were manually counted and compared to the total nuclei count using ImageJ.

For functional characterization, cortical neurons were terminally differentiated for 14 days before recording neuronal activity via the calcium dye Fluo-4AM. Time-lapse images were captured every 400 ms for 200-300 frames using a Nikon Eclipse Ti live-imaging system with a 20x Extra Long Working Distance (ELWD) objective. Neuronal activity was induced by adding a 30 mM KCl solution, prepared by diluting 1 M KCl with culture medium, to the well, making up 10% of the total volume during calcium imaging. To analyze neuronal activity, photobleaching was first corrected using the Exponential Fit method in ImageJ software. Circular regions of interest (ROIs) were drawn over neuronal cell bodies, and the mean fluorescence intensity within each ROI was measured. The intensity data were exported as .txt files and processed in PeakCaller analysis software to identify calcium peaks. The calcium trace for each ROI was exported, and the mean number of calcium peaks was plotted.

#### Cluster size analysis

Cortical neuron progenitors were plated either in a 2D conventional culture or on a bioengineered platform co-cultured with astrocytes and differentiated for 7 days. Cells were fixed, stained, and imaged using a Nikon eclipse Ti system. Cortical neuron cluster sizes were measured using DAPI stained nuclei images processed through a CellProfiler pipeline. Clusters were identified as objects based on intensity and sized filtering with diameter greater than 90 µm. The identified objects were analysed, and Max Feret diameters were extracted as representative measure of cluster sizes.

#### Aggregate count and coverage analysis

Cortical neuron progenitors were plated on a bioengineered platform using devices with aspect ratios 10, 20, and 40, with cell numbers specified in **Table 3**, either as cortical neurons alone or with astrocyte progenitors. Cells were maintained in differentiation medium for 7 days before being fixed and stained. Images were acquired using a Nikon eclipse Ti system, with a 4x objective. The number of aggregates and their percentage coverage on the plating area were analysed from DAPI-stained nuclei images. Image pre-processing was performed in ImageJ software to horizontally align linear aggregates. Each image was cropped to the same height, and the background was subtracted using a rolling size of 500 um. Aggregates were identified through intensity thresholding with the Otsu method, with a minimum size of 50 pixels (approximately 90 µm) using CellProfiler software. The detected aggregates were measured for their occupied area, and the percentage of aggregate area relative to the device area was calculated. The total area percentage was plotted against the number of aggregates.

#### Neuronal migration analysis

For migration analysis, images from the aforementioned experiment set were examined to assess neuronal migration from aggregates. Rectangular masks corresponding to the plating area of each node were created in ImageJ to define the migration area. These masks, along with DAPI- and MAP2-stained images, were analysed using CellProfiler software. DAPI and MAP2 channels were converted to binary images and identified as objects. DAPI overlapping with MAP2 were classified as MAP2-positive nuclei. MAP2-positive nuclei located outside the plating area mask and within 2 mm from the mask edge were identified, counted, and measured for distance. The count of migrated MAP2-positive nuclei was calculated per 1 mm^2^ area. The measured distances were grouped into 100 µm bins and represented as a frequency distribution to present the migration pattern.

#### Neurite directionality analysis within nodes

Neuronal nodes, with or without astrocytes, were generated on bioengineered platforms with a device aspect ratio of 20, seeded with sparsely transfected GFP cortical neuron progenitors at 16x hexagonal packing. Cells were maintained in differentiation medium for 14 days before fixation and staining. Z-stack images were acquired using an Olympus SpinSR10 SoRa spinning disk confocal microscope at 10x magnification, with a 10 µm step size. The directionality of GFP-stained neurites was analyzed using the Directionality plugin v2.3.0 in ImageJ. Images were rotated to align aggregates horizontally and grooves vertically, then cropped to a width of 1200 µm and the length of the aggregate. Image contrast was adjusted to 1% saturated pixels using the Enhance Contrast tool. Directionality analysis employed the local gradient orientation method, with 90 bins and a histogram ranging from -90° to 90°. The fraction of neurites in negative angles were combined with their corresponding positive angles, as upward and downward angles at equal deviation from 0° are equivalent. The combined fractions of neurites were plotted for each image slice. The slice labels started at 0, representing the PDMS surface, and increased with levels above the surface.

#### Density of neural aggregate analysis

Z-stack images of neuronal nodes stained with DAPI were analyzed using ImageJ software. Regions of interest (ROIs) were defined by cropping 200-µm-wide squares centered on the aggregates. Integrated intensity values were measured across the z-stack, with raw intensities corrected by subtracting background intensities measured outside the aggregates. The averaged intensity values from the middle three slices of the z-stack were calculated and plotted.

#### Neuronal activity analysis

Neuronal nodes, with or without astrocytes, were generated on bioengineered platforms with a device aspect ratio of 20, seeded with cortical neuron progenitors at 16x hexagonal packing. After 14 days of terminal differentiation, cells were stained with calcium dye Fluo-4AM and imaged using a Nikon eclipse Ti live-imaging system, with an Extra Long Working Distance (ELWD) 20x objective. Time-lapse images were acquired at 400-ms intervals for a total of 100 frames. To analyze neuronal activity, photobleaching was corrected using the Exponential Fit method in ImageJ software. Circular regions of interest (ROIs) were drawn over neuronal cell bodies, and the mean fluorescence intensity within each ROI was measured. Intensity data were exported as .txt files and processed in PeakCaller analysis software to identify calcium peaks. The mean number of calcium peaks and the mean full width at half maximum (FWHM) were calculated and converted from frame counts to time units before being plotted.

Z-stack time-lapse images of calcium activity were processed to visualize spontaneous neuronal spiking using the Temporal Colour-Code tool in ImageJ software. In the resulting images, each color represented a specific time point in the sequence, as indicated by the accompanying color code bar.

#### Analysis of neurites projecting across T-junction

Neuronal nodes were generated on bioengineered platforms with a device aspect ratio of 20, seeded with cortical and astrocytes progenitors at 8x hexagonal packing. Cells were maintained in differentiation medium for 7 days before fixation and staining. Images were acquired using a Nikon eclipse Ti widefield system, with a 4x objective. To analyse neurite projections, ROIs from β3-tubulin-stained images, positioned above T-junction were cropped using imageJ software and analyzed in CellProfiler. The CellProfiler pipeline included illumination correction to remove background and intensity measurement. For each image, the lower quartile intensity was subtracted from the total intensity, and the result was calculated as a percentage relative to the control. The average result for each well was calculated and plotted.

To determine neurite count per node, individual neurites in ROIs above the T-junction were manually counted in ImageJ, and the results were plotted.

For analysis of neurite length below the node, the distance from the end of the vertical grooves to the farthest point of neurite coverage within the T-junction was measured manually in ImageJ. Six measurements per node were averaged and expressed as a percentage of the total T-junction width.

#### Neurite tracing and distribution of neurites on the platform

Cortical circuits with two nodes were generated on a bioengineered platform, with GFP-expressing neurons in one node and non-GFP neurons in the other, both co-cultured with astrocytes progenitors, at 16x hexagonal packing. Live imaging was performed on Sir-tubulin-stained and GFP-expressing cells using a Nikon Eclipse Ti system with a 4x objective on days 2, 3, 5, 7, 12 and 14 during differentiation.

Cells were maintained in differentiation medium for 14 days before fixation and staining. Images were acquired using the Nikon Eclipse Ti system, with a 20x objective. To analyse the distribution of cortical neuron compartment localization across the platform, images were acquired from various positions, including under the node, the middle area between nodes, and both flat and grooved regions at the T-junction. ROIs with 100 µm in width were cropped from each position for analysis. The MAP2 and β3-tubulin channels were processed to generate binary images using pixel classification in the ilastik software. The area of the segmented features was subsequently quantified using Cellprofiler.

#### Directionality of axons at T-junction

Images from the aforementioned experiment set were analysed to assess the directionality of GFP-positive axons at the T-junction, using ImageJ software. To standardize orientation, images were rotated to align the T-junction grooves horizontally. ROIs with 100 µm in width were cropped from specific locations within the T-junction area. Directionality analysis was conducted using the Directionality plugin v2.3.0 with the Fourier Component method, utilizing 90 bins and a histogram range from -90° to 90°. Results were exported to an Excel spreadsheet, and the fraction of axons were grouped into three categories: vertical (-90° to -60°), intermediate (-59° to -30°), and horizontal (-29° to 0°), along with their vertical mirror reflections. The combined fractions of axons in each category were calculated and plotted.

#### Synapses analysis

Cortical neurons were plated on either ibidi glass bottom or bioengineered platforms and terminally differentiated for 21-28 days unless otherwise stated. Fixation and immunostaining were performed to label synaptic puncta (synaptophysin and PSD95) and MAP2. Imaging was performed on the Olympus SpinSR10 SoRa spinning disk confocal microscope at 60x magnification, acquiring 9 μm Z-stacks with 1 μm step size. To assess synaptic density, the colocalization of synaptophysin and PSD95 puncta within MAP2-positive dendrites was quantified (**Supplementary Figure 3A**).

**Supplementary Figure 3.**
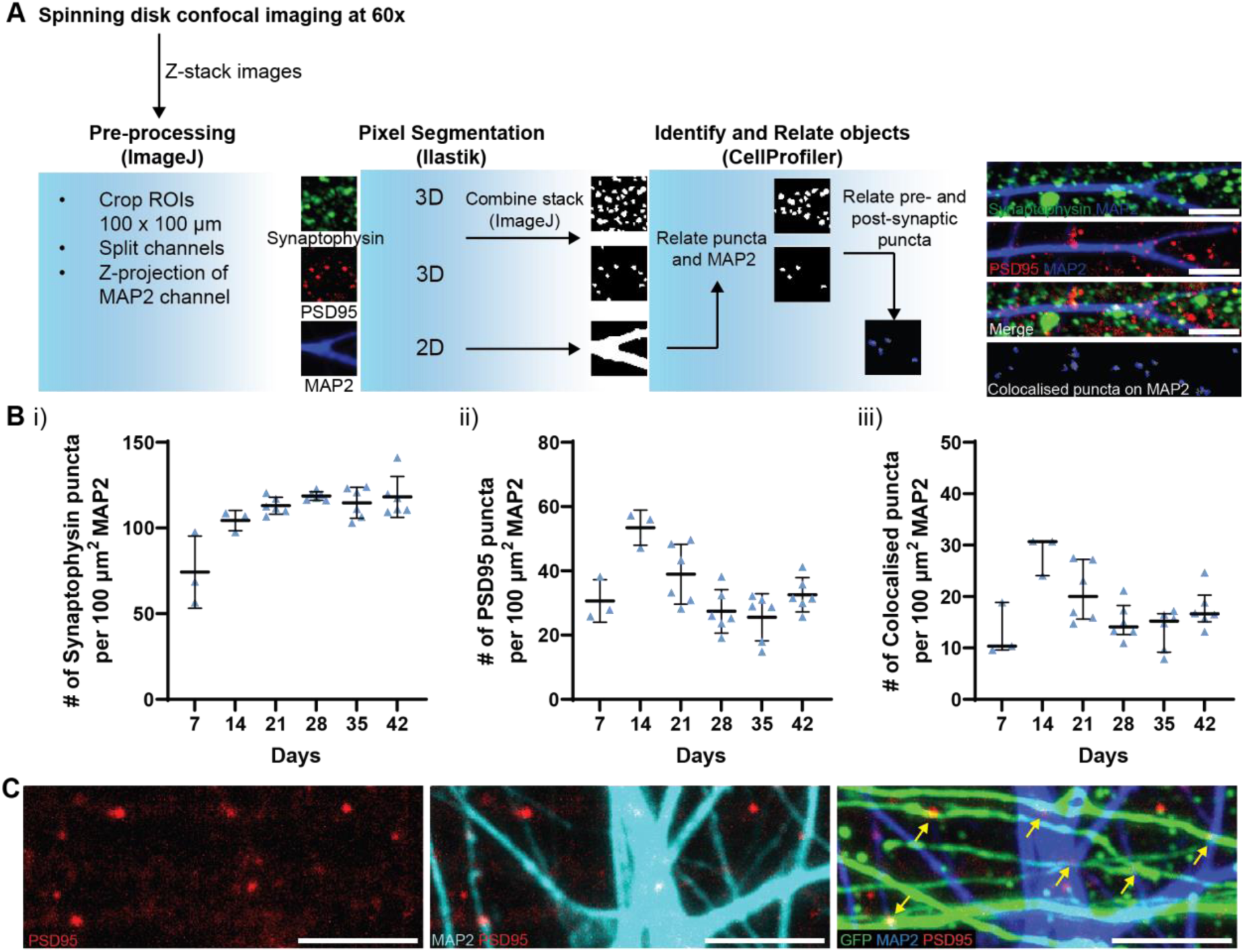
Analysis of synaptic puncta density and connectivity. A) Image analysis pipeline for synapse quantification. Z-stack images of pre-and post-synaptic protein: synaptophysin, PSD95, and neuronal structure: MAP2 were acquired from confocal microscopy with 60x magnification. Pre-processing of images was performed before object segmentation based on image pixel intensity. Each puncta object was first identified on MAP2 mask and subjected to overlapping object identification between pre-and postsynaptic puncta. Scale bar: 5 μm. B) Cortical neurons were culture on 2D ibidi glass chamber and analysed for the number of (i) pre-synaptic puncta: synaptophysin, (ii) post-synaptic puncta: PSD95, and (iii) colocalised puncta per 100 μm^2^ MAP2 positive dendrite areas, during the differentiation period from 7 to 42 days. Data points represent well averages from 1-2 independent experimental block with 1 cell line, each with 3 technical well replicates, expressed as median ± interquartile range. C) Representative images of colocalization (yellow arrow) between GFP axons from the top node and PSD95 puncta on dendrites (MAP2) of the bottom node. Scale bar: 10 μm.

**Supplementary Figure 4.**
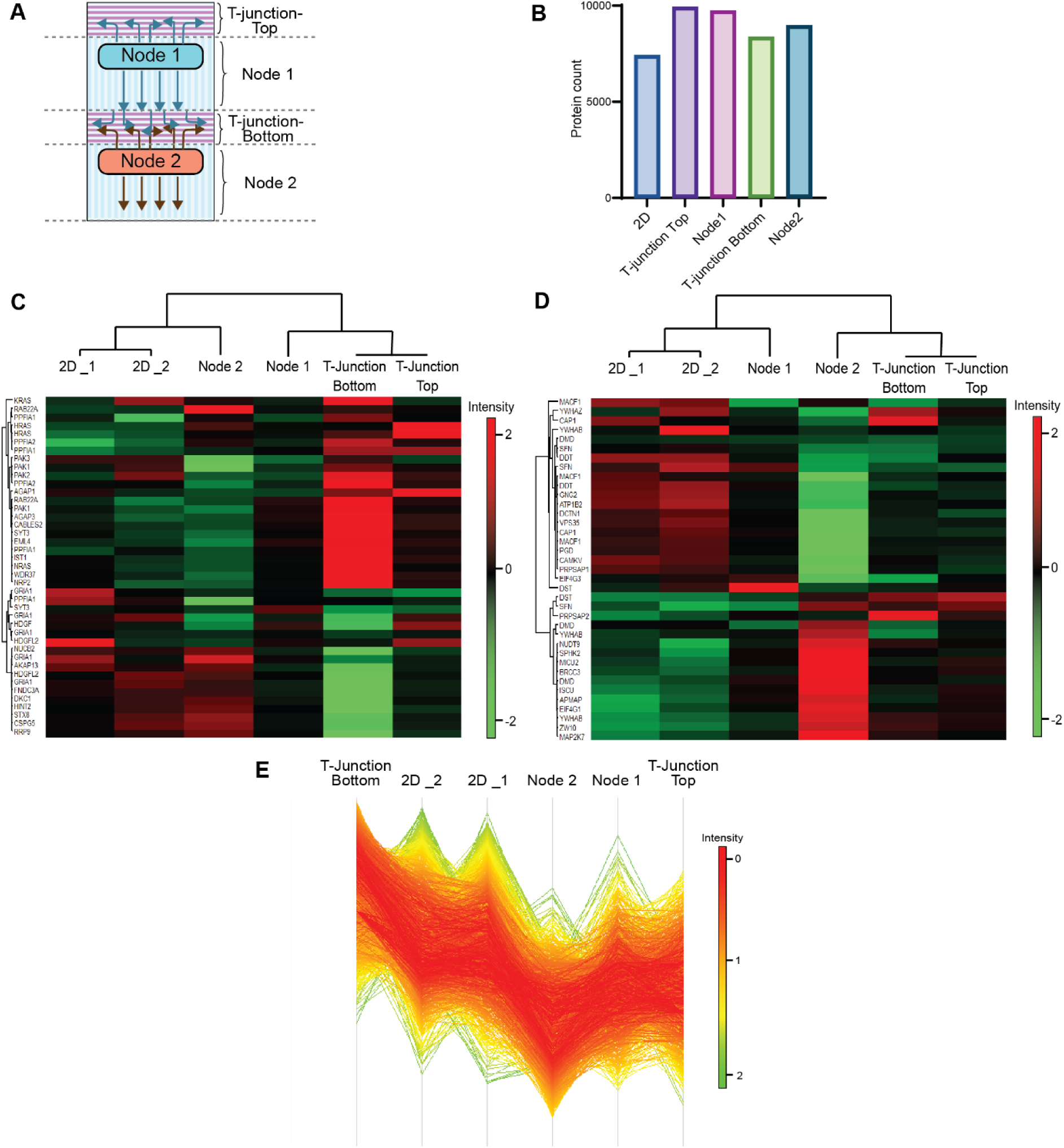
Proteomics analysis of the circuits. A) Schematic circuits on the platform, with dashed lines indicating where each fraction was cut for proteomic analysis. B) Counts of proteins analysed using Astral Orbitrap system from isolated regions of cortical networks. C) Heatmap of main contributing proteins to component 1, with samples and proteins clustered by Spearman correlation. D) Heatmap of main contributing proteins to component 2, with samples and proteins clustered by Spearman correlation. E) Density plot of the selected cluster (see Figure 7H), identifying upregulated proteins in T-junction Bottom.

Z-stack images were cropped into 100×100 µm ROIs, and each channel was spitted in ImageJ software. The MAP2 channel was projected into 2D using maximum intensity projection and smoothed with a Gaussian blur filter (σ = 1.5) to facilitate dendrite segmentation. Binary segmentations of dendrites, synaptophysin, and PSD95 were generated using the pixel classification workflow in ilastik software. Supervised training involved designating dendrites or puncta as foreground and all other elements as background, applied to 2D MAP2 images and 3D Z-stacks for synaptophysin and PSD95. Secondary antibody-only control images were considered during the training to distinguish specific synaptic puncta signals from non-specific background. After sufficient training with interactive feedback, all images were batch processed to generate segmentation output, which then subjected to batch thresholding in ImageJ to produce binary images. Binary synaptic puncta from Z-stacks were further segmented using a watershed filter, and individual spots were projected into 2D via custom ImageJ macros. All binary segmentations were processed in Cellprofiler software to identify and quantify synaptic puncta and their colocalization. Within the Cellprofiler pipeline, binary images were first converted into objects. The detected MAP2 objects were merged into a single object and related to either synaptophysin or PSD95 puncta to calculate the number of puncta per area of MAP2. The colocalization of synaptophysin and PSD95 puncta within MAP2-positive areas was subsequently quantified and reported as the number of colocalized puncta per 100 µm² of MAP2.

#### Functional connectivity

Cortical circuits on the platform were constructed using neurons genetically integrated with ChrimsonR connected to wild-type (WT) cortical neurons derived from the iPSC line KOLF2.1J. Time-lapse imaging was conducted on circuits differentiated for 21 days. Prior to imaging, cells were stained with Fluo-4AM and imaged using a Nikon Eclipse Ti live-imaging system with a 4x objective. Baseline neural activity was recorded every second for 5 minutes with 490 nm excitation. Light stimulation was applied in 100 ms pulses at 585 nm every 30 seconds during calcium imaging. Since ChrimsonR was tagged with tdTomato, cells expressing ChrimsonR were identified using 585 nm excitation. Neuronal responses to light stimulation were analyzed by drawing circular ROIs around neuron cell bodies near each node, quantifying the mean calcium intensity for each ROI over time, and processing the data with PeakCaller software to extract calcium traces.

### Statistical analysis and plotting

Data distribution was assessed using the Shapiro-Wilks or Kolmogorov-Smirnov tests. Statistical analysis was selected based on the distribution, applying parametric tests for normally distributed data and non-parametric tests for non-normally distributed data. Statistical tests and corresponding P values are indicated in the figure legends. For comparisons between two groups, the unpaired t-test was used for parametric data, and the Mann–Whitney test for non-parametric data. For comparisons across multiple groups, one-way ANOVA with Tukey’s multiple comparison test was used for parametric data, and the Kruskal–Wallis test with Dunn’s multiple comparisons for non-parametric data.

Plotting was performed using GraphPad Prism, and figures were created and assembled in Adobe Illustrator with BioRender.

### Proteomics analysis

#### Circuit construction

Cortical circuits were constructed on a bioengineered platform using neurons genetically integrated with ChrimsonR, connected to wild-type (WT) cortical neurons derived from the iPSC line KOLF2.1J. Both nodes were co-cultured with astrocytes progenitors at 16x hexagonal packing. Neurons underwent a pre-differentiation treatment one day before seeding and were further differentiated for 28 days. A control experiment was conducted using a conventional 2D 12-well plate.

#### Protein extraction

To collect circuit compartments, specific regions—including the T-junction-top, node 1, T-junction-bottom, node 2—were dissected using a surgical scalpel blade. Ten circuits per condition were pooled into a 1.5-mL Eppendorf tube and snap-frozen on dry ice. For 2D cultures, three wells from 12-well plates were pooled together into a 1.5-mL Eppendorf tube and snap-frozen on dry ice.

For protein extraction, ice-cold lysis buffer, containing 50 mM Tris, pH 7.4, 2% SDS, 10 mM Tris-(2-carboxyethyl) phosphine (TCEP), and EDTA-Free Halt protease inhibitor cocktail, was added to the samples on ice. The samples were homogenized, vortexed, and shaken at 4°C for 30 minutes. Following an additional vortexing step, PDMS pieces were removed using tweezers. The samples were then centrifuged at maximum speed for 10 minutes, and the supernatant was collected for mass spectrometry. Total protein concentration was measured using a BCA Protein Assay (ThermoFisher) following the manufacture’s guidelines.

#### Sample preparation and mass spectroscopy

For sample digestion, 10 ug of protein was processed using the SP3 methodology^31^. In short, samples were alkylated with TCEP at 56°C for 30 minutes before alkylation with Iodoacetamide for 30 minutes in the dark. Magnetic MagReSyn® Hydroxyl beads (BioResyn) were added to the samples, and proteins were washed with acetonitrile and ethanol prior to digestion. We used Trypsin / Lys-C mix (Pierce) at a 1:20 enzyme to protein ratio at 37°C overnight. The digested peptides were eluted from the beads, lyophilized using a vacuum concentrator (Thermo), and stored at -20° C.

We used 200 ng of peptides with a Vanquish Neo UHPLC System in trap and elute mode, coupled with an Orbitrap Astral Mass Spectrometer (Thermo Fisher Scientific). Peptides were loaded onto a PepMap Neo Trap Cartridge (Thermo Fisher Scientific #174500) and analyzed on a C18 EASY-Spray HPLC Column (Thermo Fisher Scientific #ES906) with a 11.8 minute gradient from 1% to 55% Buffer B (Buffer A: 0.1% formic acid in water; Buffer B: 0.08% formic acid in 80:20 acetonitrile:water, 0.7 min at 1.8 µL/min from 1% to 4% B, 0.3 min at 1.8 µL/min from 4% to 8% B, 6.7 min at 1.8 µL/min from 8% to 22.5% B, 3.7 min at 1.8 µL/min from 22.5% to 35% B, 0.4 min at 2.5 µL/min from 35% to 55% B). Eluted peptides were analyzed using data-independent acquisition mode on the mass spectrometer.

For analysis we used DIA-NN v.1.8.1 with the library free method with a human FASTA file (Uniprot, downloaded August 2023). Default parameters were used, except for “Heuristic protein inference” and the double pass neural network classifier was used.

#### Data analysis

Data was analysed using Perseus (v 2.1.3.0), following a standard workflow^32^. In short, we filtered out contaminants, then Z-score normalised using the median, and imputed missing values using a normal distribution. Graphs were produced using in-build Perseus functions and Prism. Overrepresentation analysis was performed using PANTHER^33^.

## Acknowledgments

AS whishes to acknowledge the support of BBSRC (Grant BB/W006561/1) and Motor Neurone Disease Association (Grant Serio/Apr22/885-791). PS whishes to acknowledge the support of the Ministry of Science and Technology.

## CRediT (Contributor Roles Taxonomy) Statement

*Pacharaporn Suklai* – conceptualisation, investigation, methodology, resources, validation, formal analysis, visualisation, writing – original draft, writing – review & editing

*Cathleen Hagemann* – validation, formal analysis, visualization, writing – review & editing

*Karolina Faber & Bathany Geary* – investigation, validation, formal analysis

*Taylor Minckley* – methodology, resources (generating ChrimsonR iPSC stable line and design of methodology for optogenetic system)

*Ludovica Guetta & Kelly O’toole* – resources (generating and providing iPSC-derived astrocyte progenitors)

*Rosalind Norkett & Michael J. Devine* – resources (dissection and preparation of mouse cortical neurons)

*Andrea Serio* – conceptualisation, supervision, funding acquisition, project administration, visualisation, writing – original draft, writing – review & editing

